# TIR-catalyzed ADP-ribosylation reactions produce signaling molecules for plant immunity

**DOI:** 10.1101/2022.05.02.490369

**Authors:** Aolin Jia, Shijia Huang, Wen Song, Junli Wang, Yonggang Meng, Yue Sun, Lina Xu, Henriette Laessle, Jan Jirschitzka, Jiao Hou, Tiantian Zhang, Wenquan Yu, Giuliana Hessler, Ertong Li, Shoucai Ma, Dongli Yu, Jan Gebauer, Ulrich Baumann, Xiaohui Liu, Zhifu Han, Junbiao Chang, Jane E. Parker, Jijie Chai

**Affiliations:** Beijing Advanced Innovation Center for Structural Biology, Tsinghua-Peking Joint Center for Life Sciences, Center for Plant Biology, School of Life Sciences, Tsinghua University, Beijing 100084, China; Institute of Biochemistry, University of Cologne, Cologne 50674, Germany; Max Planck Institute for Plant Breeding Research, Department of Plant-Microbe Interactions, Cologne 50829, Germany; Henan Key Laboratory of Organic Functional Molecules and Drug Innovation, Henan Normal University, Xinxiang 453007, China; School of Pharmaceutical Sciences, Zhengzhou University, Zhengzhou 450001, China; College of Chemistry, Zhengzhou University, Zhengzhou 450001, China; National Protein Science Facility, Tsinghua university, Beijing 100084, China

## Abstract

Plant pathogen-activated immune signaling by nucleotide-binding leucine-rich repeat (NLR) receptors with an N-terminal Toll/Interleukin-1 receptor (TIR) domain converges on Enhanced Disease Susceptibility 1 (EDS1) and its direct partners Phytoalexin Deficient 4 (PAD4) or Senescence-Associated Gene 101 (SAG101). TIR-encoded NADases produce signaling molecules to promote exclusive EDS1-PAD4 and EDS1-SAG101 interactions with helper NLR sub-classes. Here we show that TIR-containing proteins catalyze adenosine diphosphate (ADP)-ribosylation of adenosine triphosphate (ATP) and ADP ribose (ADPR) via ADPR polymerase-like and NADase activity, forming ADP-ribosylated ATP (ADPr-ATP) and ADPr-ADPR (di-ADPR), respectively. Specific binding of di-ADPR or ADPr-ATP allosterically promotes EDS1-SAG101 interaction with helper NLR N requirement gene 1A (NRG1A) *in vitro* and *in planta*. Our data reveal an enzymatic activity of TIRs that enables specific activation of the EDS1-SAG101-NRG1 immunity branch.

---

Cell-surface and intracellular immune receptors detect pathogen disturbance and initiate signaling cascades leading to host induced defenses (*1, 2*). The Toll-Interleukin-1 receptor (TIR) domain is an immune signaling module in many species (*2*). Bacterial and plant TIRs, and human sterile alpha and TIR motif containing 1 (hSARM1) are NAD^+^-hydrolyzing enzymes (*3–5*). In plants, intracellular nucleotide-binding/leucine-rich repeat (NLR) receptors recognize pathogen-delivered effectors to confer effector-triggered immunity (ETI) (*6*). ETI transcriptionally boosts generally weaker basal immunity mediated by cell-surface pattern-recognition receptors conferring pattern-triggered immunity (PTI), often leading to localized host cell death (*7, 8*). A major pathogen-sensing NLR sub-class (called TIR-NLRs or TNLs) has N-terminal TIR signaling domains (*2, 5*). Also, TIR-domain proteins which lack leucine-rich repeats (TIRs) contribute to ETI and basal immunity responses (*5, 9–11*). The signaling role of TNLs and TIRs depends on their NADase activity (*3, 4, 12*), which is promoted by effector-induced TNL oligomerization (*12, 13*) or TIR self-association (*3, 4, 14*).

Plant immunity mediated by TNLs and TIRs requires the conserved lipase-like protein EDS1 and its sequence-related direct partners, PAD4 or SAG101 (*15–17*), and two sub-families of coiled-coil (CC)-like N-terminal domain helper NLRs, ADR1s and NRG1s (*16, 18–22*). Studies in *Arabidopsis* and *Nicotiana benthamiana* (*Nb*-tobacco) revealed that EDS1-SAG101 cooperates with NRG1-type helper NLRs as a TNL ETI-induced module promoting host cell death (*16, 23*). Functional cooperation and pathogen-induced interaction between the *Arabidopsis* EDS1-PAD4 dimer and ADR1-type helper NLRs, by contrast, is associated with boosted basal immunity and restriction of pathogen growth (*16, 18, 23-26*). In *Arabidopsis*, these two modules are partially redundant in their functions, but they also display preferences for specific immune outputs (*27*).

TIR-catalyzed small molecules induced the assembly of *Arabidopsis* EDS1-SAG101-NRG1 and EDS1-PAD4-ADR1 complexes in insect cells (*28*). TNL and TIR-generated 2’-(5’’-phosphoribosyl)-5’-adenosine mono-/di-phosphate (pRib-ADP/AMP) efficiently induced EDS1-PAD4 association with ADR1-L1 but displayed only weak activity in promoting EDS1-SAG101 interaction with NRG1A (*28*). These data and the EDS1-SAG101-NRG1 signaling preferences *in vivo* suggested that other, yet unidentified, TNL/TIR-catalyzed small molecules underlie signaling via this immune complex. Here we show that TIR domains catalyze an ADP ribose-transfer reaction using NAD^+^ or NAD^+^ with ATP as substrates to produce, respectively, ADP-ribosylated ADPR (di-ADPR) and ADP-ribsylated ATP (ADPr-ATP). Like pRib-ADP/AMP (*28*), di-ADPR and ADPr-ATP can function as second messengers to induce TNL signaling. The latter two molecules specifically bind to and induce *Arabidopsis* EDS1-SAG101 interaction with NRG1A, establishing EDS1-SAG101 as a receptor complex for di-ADPR and ADPr-ATP. Structural analyses reveal specific recognition mechanisms for di-ADPR and ADPr-ATP by EDS1-SAG101. Our data show that different small molecules catalyzed by TIR proteins selectively potentiate two distinctive EDS1-mediated signaling complexes, and shed light on the catalytic mechanism underlying production of these small molecules.

## Identification of a small molecule specifically activating EDS1-SAG101-NRG1

pRib-AMP/ADP displayed weak activity in inducing the EDS1-SAG101 interaction with NRG1A (*28*), suggesting that other TIR-catalyzed small molecules promote this interaction more efficiently. When co-expressed with the *Arabidopsis* TNL RPP1 and its cognate *Hyaloperonospora arabidopsidis* effector ATR1 (*29*), EDS1-SAG101 was co-purified with N-terminally GST-fused NRG1A (fig. S1A) (*28*). NRG1A binding of EDS1-SAG101 was suppressed by mutation of RPP1 catalytic residue Glu158, suggesting that EDS1-SAG101 interaction with NRG1A is conferred by RPP1-catalyzed products. We denatured the insect cell-expressed EDS1-SAG101 protein and extracted EDS1-SAG101-bound small molecule(s). While separately purified apo-EDS1-SAG101 failed to interact with NRG1A, the complex gained NRG1A-but not ADR1-L1-binding activity when incubated with the extractant (fig. S1B). Liquid chromatography and high-resolution mass spectrometry (LC-HRMS) analysis of the extractant revealed a chemical species with mass-to-charge ratio (m/z) 1047.0530 (z=1^-^) (Fig. 1A). Additionally, a trace amount of a product with molecular weight matching that of pRib-ADP was detected (fig. S1C), supporting weak pRib-ADP activity in promoting EDS1-SAG101 association with NRG1A (*28*).

**Fig. 1.**
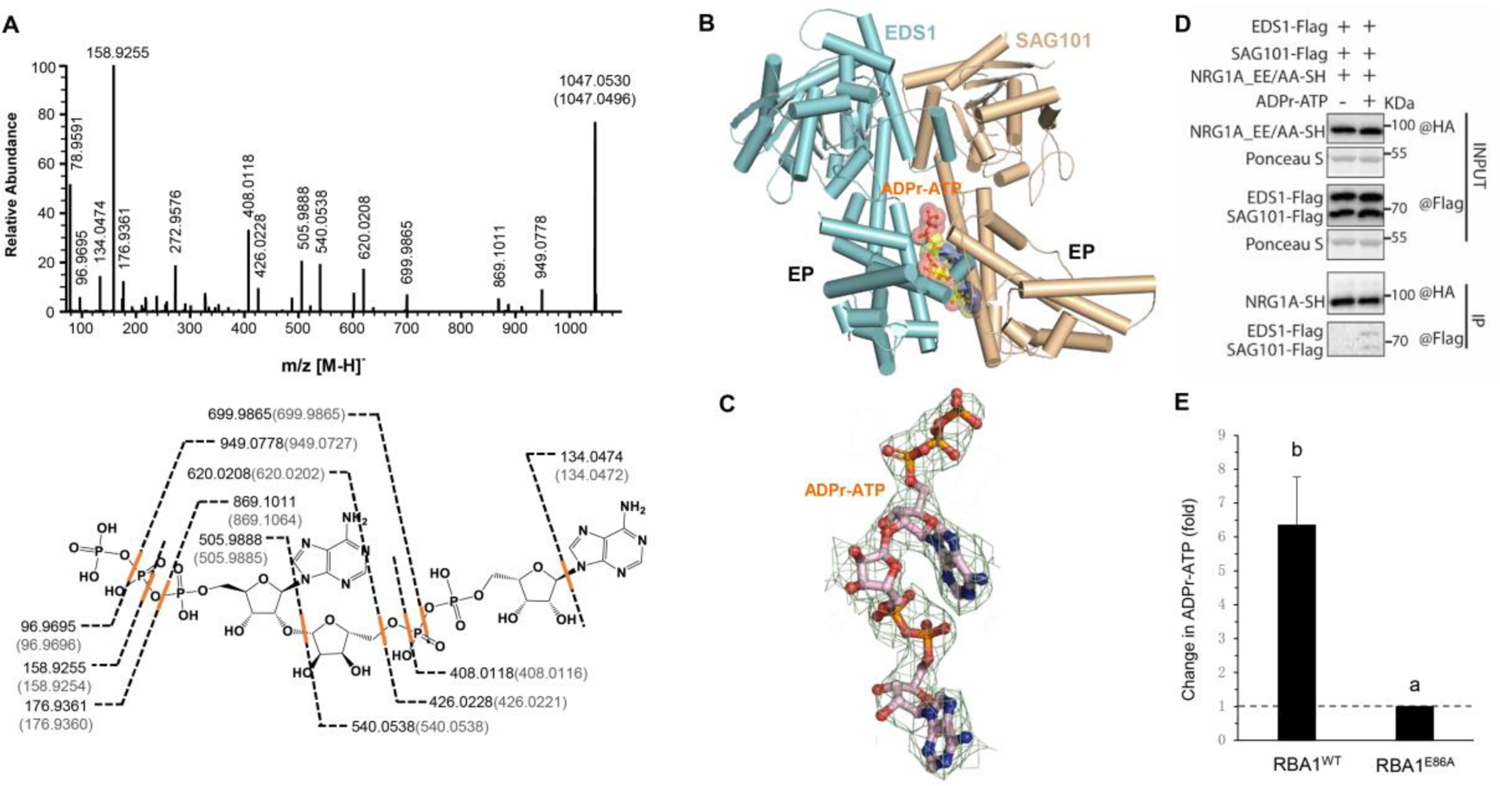
Identification of a TIR-catalyzed small molecule activating EDS1-SAG101. (**A**) Mass analysis of the small molecule bound to EDS1-SAG101. Top: MS/MS spectra after collision induced dissociation (CID) fragmentation of the ion with m/z = 1047.0530 (z =1^-^). Bottom: Scheme showing the proposed routes of formation of various ions shown above. Numbers in parentheses indicate theoretical molecular weights of proposed ions. (**B**) Cryo-EM structure of EDS1-SAG101 bound by ADPr-ATP. ADPr-ATP bound between EDS1 and SAG101 EP domains is shown in transparent sphere. EDS1 and SAG101 are shown in cartoon and ADPr-ATP in stick. (**C**) Cryo-EM density around the bound ADPr-ATP (in stick). (**D**) ADPr-ATP induces EDS1-SAG101 interaction with NRG1A after protein expression in *Nb* tobacco. ADPr-ATP previously extracted from EDS1-SAG101 was incubated with a lysate of *Nb-*tobacco quadruple mutant *eds1pad4sag101ab* (*Nb-epss*) expressing EDS1-FLAG, SAG101-FLAG and NRG1A^E14A/E27A^-SH at 4°C for 1 h. Protein-protein interaction was detected by Co-IP with antibodies as indicated. (**E**) Relative change in ADPr-ATP accumulation in *Nb-epss* plants expressing wild type RBA1 (RBA1^WT^) or RBA1^E86A^. The statistics are based on LC-MS results of three replicates. The amount of ADPr-ATP in RBA1^E86A^-expressing *Nb-epss* was normalized to 1.0. Significance calculated with Tukey’s HSD test (n=3, α=0.05, different letters indicate significant differences). *Nb-epss* leaves expressing RBA1^WT^ or the catalytic mutant *RBA1*^E86A^ and insect cells expressing apo-EDS1-SAG101 were mixed and co-sonicated. The apo-EDS1-SAG101 complex in the lysate was purified through affinity chromatography. Multiple reaction monitoring (MRM) analyses were performed for ADPr-ATP bound in the purified apo-EDS1-SAG101 complex.

To identify the chemical species with m/z 1047.0530 (z=1^-^), we solved a cryo-EM structure of EDS1-SAG101 purified after co-expression with RPP1 and ATR1 in insect cells (Fig. 1B, fig. S2 and Table S1). A cryo-EM density not occupied by EDS1-SAG101 is clearly discerned at the interface between the C-terminal EP domains of EDS1 and SAG101 (Fig. 1C), corresponding to where pRib-ADP binds in EDS1-PAD4 (fig. S3A) (*28*). pRib-ADP fitted well into half of the unoccupied cryo-EM density (fig. S3B), but two patches of cryo-EM density remained unfilled following pRib-ADP modeling (fig. S3B). The smaller patch, which connects to the terminal (β)-phosphate group of the modeled pRib-ADP, likely results from a phosphate group. The other patch is much larger and accommodates the AMP molecule. The phosphate group, pRib-ADP and AMP are subgroups of ADPr-ATP, with a calculated molecular mass of 1048.0496 matching 1048.0530 from the LC-HRMS analysis (Fig. 1A, top). Furthermore, prominent ions with m/z values expected for product ions of ADPr-ATP were observed in the high-resolution MS/MS spectrum (Fig. 1A, bottom).

We investigated whether ADPr-ATP extracted and purified as described above induces EDS1-SAG101-NRG1A complex when expressed in *Nb* tobacco. *EDS1, SAG101* and an *NRG1A* E14A/E27A cell death-inactive variant (*23, 30, 31*) were transiently co-expressed in an *Nb* tobacco *epss* line lacking all EDS1-family proteins (*Nb-epss*) (*16*). ADPr-ATP was incubated with the *Nb-epss* tissue lysate. Incubation ADPr-ATP resulted in an interaction between EDS1-SAG101 and NRG1A which was not detected in the absence of ADPr-ATP (Fig. 1D). These results indicate that ADPr-ATP is sufficient to promote EDS1-SAG101 association with NRG1A.

We next employed the *Nb-epss* transient assay to investigate whether a TIR protein catalyzes accumulation of ADPr-ATP in plants. *Arabidopsis* RBA1 (*9*) was used for this because of its *EDS1*-dependent cell death activity in *Nb* tobacco (*28*). *Nb*-*epss* leaves expressing RBA1 were co-sonicated with insect cells expressing EDS1 and SAG101, and LC-MS was used to detect small molecules captured by purified EDS1-SAG101. As shown in Fig. 1E, plants expressing wild type RBA1 but not an RBA1 catalytic mutant (E86A) produced strong ADPr-ATP signals in the assay. These data show that TIR-catalyzed ADPr-ATP produced in plant tissues binds to EDS1-SAG101.

## Recognition mechanism of ADPr-ATP by EDS1-SAG101

ADPr-ATP adopts an elongated conformation binding to a positively charged pocket between EDS1 and SAG101 EP domains (Fig. 2A). The pRib-ADP moiety in ADPr-ATP forms a large network of hydrogen bonding and polar interactions with EDS1-SAG101 (Fig. 2B) that are conserved in the structure of pRib-ADP-bound EDS1-PAD4 (fig. S3C). The remaining portions of ADPr-ATP also establish extensive contacts with EDS1-SAG101. The terminal nucleobase adenine hydrophobically packs against SAG101 Met384 and Leu438 at one side and EDS1 His476 at the other side of the pocket (Fig. 2B). The terminal phosphate group is solvent-exposed and only makes van der Waals contact with SAG101 Lys371. Sequence alignment showed that the ADPr-ATP-interacting residues of EDS1 (fig. S4A) and SAG101 (fig. S4B) are conserved among dicot plant species.

**Fig. 2.**
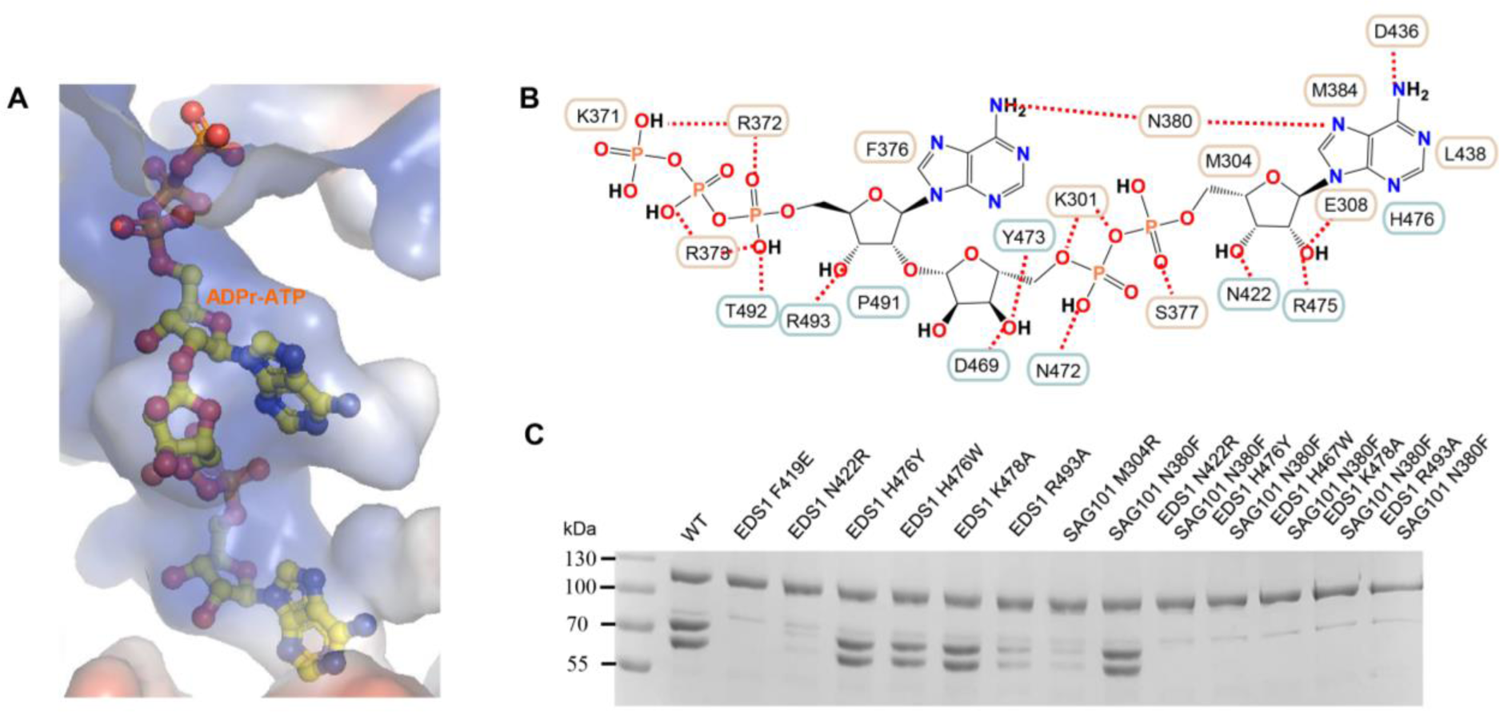
ADPr-ATP specific recognition by EDS1-SAG101. (**A**) Close-up view of the ADPr-ATP-binding pocket of EDS1-SAG101. EDS1-SAG101 is shown in electrostatic surface and ADPr-ATP in stick. Blue, red and white represent positive, negative and neutral surfaces, respectively. (**B**) Diagram showing interaction between EDS1-SAG101 and ADPr-ATP. Cyan and purple frames represent residues from EDS1 and SAG101, respectively. Red dashed lines represent polar interactions. (**C**) Analysis of EDS1-SAG101 ADPr-ATP binding pocket mutants for induced association with NRG1A. N-terminally GST-NRG1A was co-expressed with non-tagged EDS1 and C-terminally strep-tagged SAG101 (wild type or mutant forms as indicated), RPP1 and 10xHis-tagged ATR1 in insect cells. GST-NRG1A was purified using Glutathione Sepharose 4B (GS4B) beads. GS4B-bound proteins were eluted, separated by SDS-PAGE and detected by Coomassie Brilliant Blue staining.

To verify the ADPr-ATP activated EDS1-SAG101 cryo-EM structure, we assessed the impact of mutations at the ADPr-ATP-binding pocket on EDS1-SAG101 interaction with NRG1A using the pulldown assay described above. Substitution of SAG101 Met304 with its PAD4 equivalent Arg314 substantially weakened EDS1-SAG101 interaction with NRG1A (Fig. 2C), explaining why ADPr-ATP was barely active in stimulating EDS1-PAD4 interaction with ADR1-L1 (fig. S1B). In *Arabidopsis*, SAG101 Met304 and PAD4 Arg314 were essential for signaling through their respective branches and induced association with corresponding helper NLRs (*32*). In contrast, interaction was only slightly impacted by substitution of SAG101 Asn380 with its PAD4 equivalent Phe387 (Fig. 2C). Supporting a specific role for EDS1 His476 in ADPr-ATP recognition, H476Y substantially suppressed AvrRps4-induced cell death but had little effect on AvrRps4-induced immunity in *Arabidopsis* (*16*). While EDS1 H476Y mutation did not disable EDS1-SAG101 interaction with NRG1A (Fig. 2C), combination of SAG101 N380F and EDS1 H476Y resulted in loss of NRG1A binding activity of EDS1-SAG101. An R493A mutation only reduced induced EDS1-SAG101 interaction with NRG1A (Fig. 2C), in contrast with a loss of pRib-ADP-induced EDS1-PAD4 interaction with ADR1-L1 (*28*). This would explain why an EDS1 R493A variant disturbed mainly EDS1-PAD4 mediated pathogen resistance *in vivo* (*16, 25*). The difference might be due to more extensive contacts between ADPr-ATP and EDS1-SAG101, rendering binding of the small molecule less sensitive to mutation at this position.

## Selective EDS1-SAG101 activation mechanisms by ADPr-ATP

ADPr-ATP specifically promotes EDS1-SAG101 interaction with NRG1A (fig. S1B). By comparison, pRib-ADP induces EDS1 heterodimer interactions with ADR1-L1 and NRG1A, although with different efficiency (*28*). pRib-ADP and ADPr-ATP bind to a similar position in the EDS1-PAD4 and EDS1-SAG101 complexes (fig. S3A). The pRib-ADP-contacting residues of PAD4 are largely conserved in SAG101 (fig. S3C), whereas ADPR-interacting residues of SAG101 are largely variable in PAD4 (fig. S5). Notably, substitutions of SAG101 Met304 and Asn380 with their bulkier equivalents PAD4 Arg314 and Phe387 result in occlusion of the ADPR-binding site in EDS1-PAD4 (Fig. 3A). The different rotamers of PAD4 Gln318 and its counterpart SAG101 Glu308 further sterically block ADPr-ATP from being recognized by EDS1-PAD4.

**Fig. 3.**
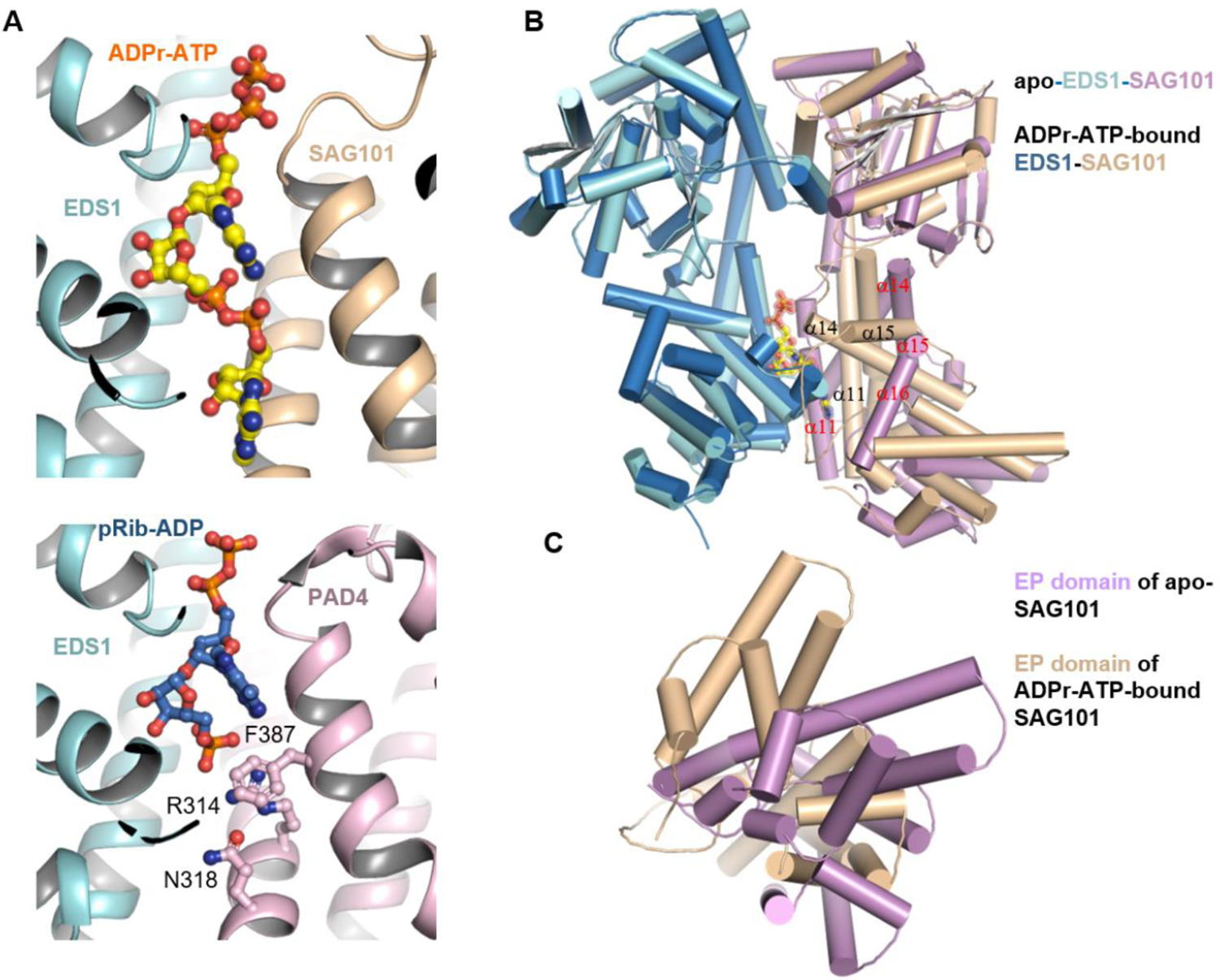
ADPr-ATP selection and activation mechanisms of EDS1-SAG101. (**A**) Structural alignment of EDS1-SAG101 (top) and EDS1-PAD4 (bottom) around the ligand-binding sites. EDS1 (cyan), SAG101 (light orange) and PAD4 (pink) are shown in cartoon, and ADPr-ATP (yellow) and pRib-ADP (blue) in stick. Three PAD4 residues Phe387, Arg314 and Asn318 not conserved in SAG101 are shown. (**B**) Structural alignment of apo-EDS1-SAG101 and ADPr-ATP-bound EDS1-SAG101. Color codes are indicated. Secondary structural elements of SAG101 around the ADPr-ATP-binding site are labeled. (**C**) Bottom view of the EP domain structural alignment between apo-EDS1-SAG101 and ADPr-ATP-bound EDS1-SAG101 shown in (B).

Superposition of an apo-EDS1-SAG101 crystal structure (*15*) with the cryo-EM structure of ADPr-ATP-bound EDS-SAG101 revealed that the EDS1 conformation is nearly identical in the two forms of EDS1-SAG101 (Fig. 3B). There is a striking structural difference in the SAG101 EP domain, while only a slight conformational change in the SAG101 lipase-like domain upon ADPr-ATP binding (Fig. 3B). Compared to the apo-form of SAG101, the EP domain in ADPr-ATP bound SAG101 rotates anti-clockwise about 15 degrees (viewed from the bottom) (Fig. 3C). This causes complete melting of α16 and shifts of the N-terminal sides of helices α14 and α15 towards EDS1, whereas SAG101 α11 shifts away from EDS1 about 4.0 Å. These structural analyses indicate that ADPr-ATP binding triggers conformational changes in the SAG101 EP domain which enable EDS1-SAG101 interaction with NRG1A. Thus, ADPr-ATP and pRib-ADP (*28*) employ a conserved allosteric mechanism to activate EDS1-SAG101 and EDS1-PAD4, respectively.

## TIR-catalyzed di-ADPR induces EDS1-SAG101 interaction with NRG1A

Expression of RBA1 resulted in accumulation of ADPr-ATP in *Nb* tobacco (Fig. 1E). We next investigated whether a purified TIR protein catalyzes production of ADPr-ATP when NAD^+^ is used as a substrate. RPS4 TIR domain (residues 1-236, RPS4-TIR) which causes HR-like cell death when expressed *in planta* (*33*), was used because it was easily purified. As shown in Fig. 4A, incubation of EDS1-SAG101 with wild type RPS4-TIR but not its catalytic mutant E88A in the presence of NAD^+^ induced EDS1-SAG101 interaction with NRG1A. By contrast, EDS1-SAG101 displayed no NRG1A-binding activity in the absence of NAD^+^. These results suggest that RPS4-TIR catalyzed ADPr-ATP production supports EDS1-SAG101 interaction with NRG1A. However, LC-HRMS analysis of extracted chemical compounds from the EDS1-SAG101 protein incubated with RPS4-TIR and NAD^+^ revealed a strong peak at m/z 1099.1276 (z=1^-^) (Fig. 4B) instead of the predicted m/z 1047.0530 (z=1^-^) of ADPr-ATP. Given its EDS1-SAG101-binding activity, the unknown substance with m/z 1099.1276 (z=1^-^) should be structurally related to ADPr-ATP. One such candidate is di-ADPR with molecular weight (1100.1255) which matches the 1100.1276 peak measured by LC-HRMS. Supporting this assignment, prominent ions with m/z values expected for product ions of di-ADPR were observed in the high-resolution MS/MS spectrum (Fig. 4C).

**Fig. 4.**
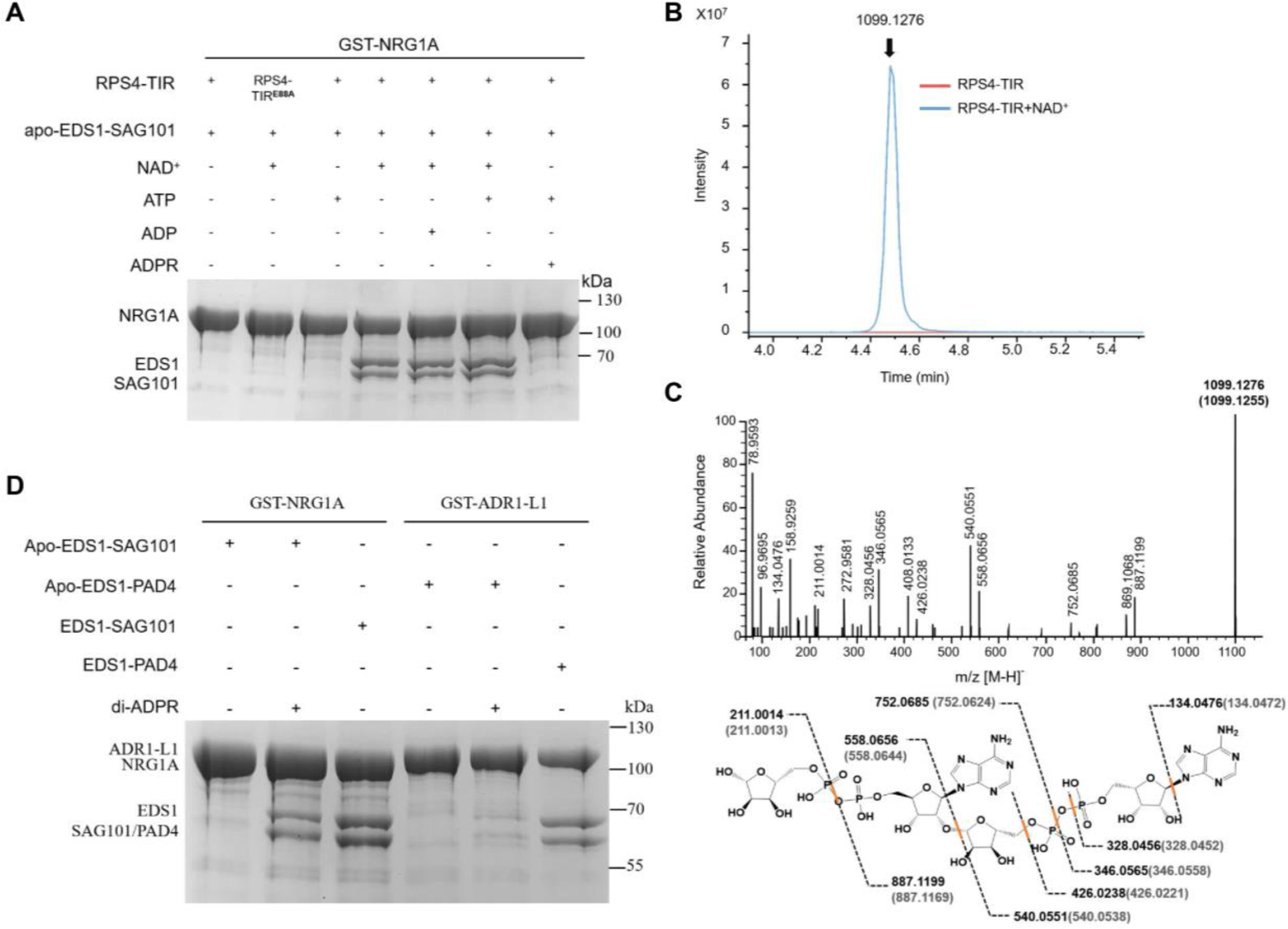
TIR-catalyzed di-ADPR induces EDS1-SAG101 interaction with NRG1A. (**A**) RPS4 TIR catalyzes different substrates to produce small molecules inducing EDS1-SAG101 interaction with NRG1A. Reaction products of RPS4 TIR domain (residues 1-236) protein with substrates indicated were incubated with GS4B beads bound by GST-tagged NRG1A preincubated with excess apo-EDS1-SAG101 on ice for 30 min. After washing, proteins bound in GS4B beads were eluted, analyzed by SDS-PAGE and detected by Coomassie Brilliant Blue staining. (**B**) LC-HRMS chromatograms of EDS1-SAG101-captured small molecules from RPS4-TIR catalyzed products with NAD^+^ as the substrate. Purified RPS4-TIR protein was incubated with NAD^+^ at room temperature for 16 h. Products of the reaction were captured by apo-EDS1-SAG101 and the captured small molecules were extracted by denaturing the complex with 80% methanol (v/v) and then subjected to LC-HRMS. (**C**) Mass analysis of small molecules in (B) Top: MS/MS spectra after collision induced dissociation (CID) fragmentation of the ion with m/z = 1099.1276 (z =1^-^). Bottom: Scheme showing proposed routes of formation of various ions shown above. Numbers in parentheses indicate theoretical molecular weights of proposed ions. (**D**) A synthetic di-ADPR enanthiomer induces EDS1-SAG101 interaction with NRG1A. The synthetic compound was incubated with GS4B beads bound by GST-tagged NRG1A/ADR1-L1 preincubated with excess apo-EDS1-SAG101/ apo-EDS1-PAD4 on ice for 30 min. After washing, proteins bound in GS4B beads were eluted, analyzed by SDS-PAGE and detected by Coomassie Brilliant Blue staining.

The activity of ADPr-ATP and di-ADPR in inducing EDS1-SAG101 interaction with NRG1A suggests that the terminal ribose sugar ring of di-ADPR or the γ-phosphate of ADPr-ATP is not essential for EDS1-SAG101 binding. This is consistent with the observation that the γ-phosphate is solvent-exposed in the cryo-EM structure of EDS1-SAG101 (Fig. 1B). Further supporting this conclusion, a chemically synthesized di-ADPR enantiomer with the configuration of C-1’’ from the terminal ribose sugar ring inverted (fig. S7) induced the EDS1 heterodimer interaction with NRG1A (Fig. 4D). By contrast, the synthetic compound had little activity of potentiating EDS1-PAD4 binding to ADR1-L1 in the assay.

## Mechanism of TIR-catalyzed production of bioactive small molecules

TIR-catalyzed production of ADPr-ATP seemingly results from condensation between ATP and ADPR. However, incubation with RPS4-TIR, ATP and ADPR failed to promote EDS1-SAG101 interaction with NRG1A (Fig. 4A). Thus, neither ADPr-ATP nor di-ADPR was produced in the reaction and ADPR is unlikely to be the RPS4-TIR sole substrate for di-ADPR production. Given that NADase activity is required for TIR-induced EDS1-SAG101 interaction with NRG1A (Fig. 4A and fig. S1A), we reasoned that formation of the 1”-2′ glycoside bond in ADPr-ATP and di-ADPR involves NAD^+^ hydrolysis. Indeed, incubation of an equal molar ratio of ATP and NAD^+^ with RPS4-TIR conferred EDS1-SAG101 NRG1A-binding activity (Fig. 4A). LC-HRMS showed that both di-ADPR and ADPr-ATP were produced in this reaction with di-ADPR being the major product (fig. S6A). However, ADPr-ATP was predominantly captured by EDS1-SAG101 (fig. S6B), suggesting a higher EDS1-SAG101 affinity for ADPr-ATP than di-ADPR.

A crystal structure of RPP1-TIR (residues 83-254) solved in the presence of NAD^+^ revealed that the TIR domain forms a tetramer which is nearly identical to that in the cryo-EM structure of the RPP1 resistosome (fig. S8 and Table S2) (*12*). Well-resolved electron densities at the two asymmetric dimer interfaces (Fig. 5A) where the ATP molecules are bound in the RPP1 (*12*) and ROQ1 (*13*) resistosomes (fig. S8A), match well with ADPR, indicating that the crystal structure represents an RPP1 TIR-product complex. C1’’ in the ribose sugar ring of ADPR faces the catalytic residue Glu158 (Fig. 5B), suggesting that this residue is likely important for catalyzing nicotinamide release from NAD^+^. ADPR binding to the asymmetric RPP1 TIR dimer is remarkably similar to that of a substrate mimic to the active site of hSARM1 (fig. S8B) (*34*). These data suggest that formation of the 1”-2′ glycoside bond in ADPr-ATP involves nucleophilic attack on the C1’’ of the TIR-activated NAD^+^ by the 2’-OH of ATP, resulting in ADP-ribosylation of the in-coming ATP molecule. ADPR is known to be a product of TIR-catalyzed NAD^+^ hydrolysis (*3, 4*). Thus, di-ADPR is produced via ADP-ribosylation of ADPR.

**Fig. 5.**
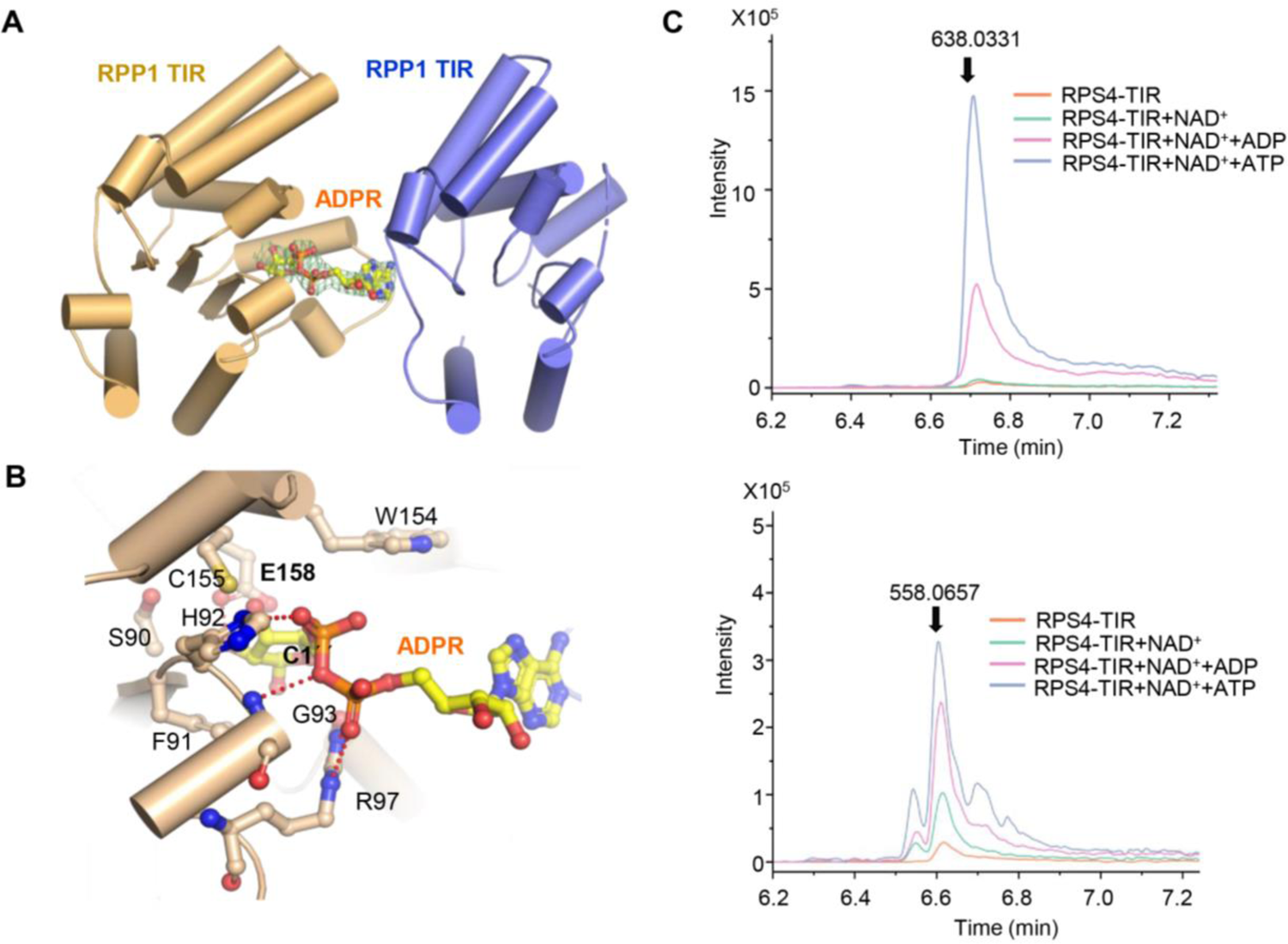
Mechanism for TIR-catalyzed production of ADPr-ATP, di-ADPR and pRib-AMP/ADP. (**A**) An asymmetric RPP1 TIR dimer with ADPR bound. Omit electron density (2*Fo*-*Fc*) contoured at 1.3 sigma surround ADPR (in stick). (**B**) Detailed interactions between the ribose sugar ring and one RPP1 TIR. Red dashed lines represent polar interactions. the C-1’’ atom is labeled. (**C**) LC-HRMS chromatograms of the apo-EDS1-PAD4-captured pRib-ADP (top) and pRib-AMP (bottom) from RPS4-TIR catalyzed products. Different colors represent different substrate of RPS4-TIR. Purified RPS4-TIR protein was incubated with substrates indicated at room temperature for 16 h. Products of each reaction were captured by apo-EDS1-PAD4 and the captured small molecules were extracted by denaturing the complex with 80% methanol (v/v) and subjected to LC-HRMS analyses.

Our previous study showed that TIR proteins can hydrolyze ATP/GTP (*35*). We therefore tested whether pRib-AMP/ADP (which efficiently activate EDS1-PAD4) (*28*) can be produced through RPS4-TIR hydrolysis of ADPr-ATP or di-ADPR. NAD^+^, NAD^+^+ATP or NAD^+^+ADP were incubated with bacteria-purified RPS4-TIR and insect cell-purified EDS1-PAD4 was used to capture the products of each reaction. The EDS1-PAD4-captured molecules were detected by LC-HRMS. Both pRib-ADP and pRib-AMP were captured by EDS1-PAD4 when ATP+NAD^+^ were the substrates (Fig. 5C). By comparison, much lower pRib-ADP amounts were captured when NAD^+^ alone was the substrate. Co-incubation of NAD^+^ with ADP notably promoted production of pRib-ADP/AMP. These data point to a two-step reaction in which RPS4-TIR first generates ADPr-ADP via ADP-ribosylation of ADP and then hydrolyzes the product to form pRib-ADP/AMP (fig. S9).

## Discussion

We discover here and in a related study (*28*) a broadly conserved allosteric activation mechanism for activation of EDS1-PAD4 and EDS1-SAG101 by TIR-catalyzed small molecules. The induced conformational change in the two EDS1 heterodimers promotes direct association with co-functioning helper NLRs, presumably activating their Ca^2+^-permeable ion channel activity (*30*). Despite the shared biochemical activity, resulting EDS1-PAD4-ADR1 and EDS1-SAG101-NRG1 signaling complexes display preferences for specific immune outputs (*16, 23, 32*). This can be partly reconciled by the existence of multiple TIR-catalyzed signaling molecules with overlapping and distinct specificity for interactions with the two EDS1 heterodimers (*28*) (Fig. 4A). Amino acids that influence selection of these small molecules by EDS1 heterodimers (Figs. 2 and 3A) and immunity functions *in vivo* (*16, 23, 25, 32*) are located at similar EDS1-PAD4 and EDS1-SAG101 EP-domain surfaces. Notably, an EDS1 mutation (H476Y) that perturbed the ligand binding site of EDS1-SAG101 but not EDS1-PAD4 (Figs. 2B and 3A) abolished host cell death but not resistance to bacteria in *Arabidopsis* (*16*). Thus, specific immune functions of the two EDS1 heterodimers appear to be conferred by their recognized ligands. This is consistent with the observation that overexpression of EDS1 with PAD4 leads to increased Arabidopsis pathogen resistance but not host cell death (*24*). The strong activity of pRib-AMP in promoting EDS1-PAD4 interaction with ADR1 but a very low capacity for promoting EDS1-SAG101 interaction with NRG1A (*28*) suggests that the TIR product might induce effective plant immunity without cell death. The identified TIR-catalyzed signaling molecules are produced with unequal molar ratios (Fig 5C and fig. S6A), probably due to differences in their stability (*28*) and production efficiency. In plant tissues, TIR-catalyzed small molecules might be subject to temporal and spatial regulation to ensure that one particular set accumulates. Such regulation might be an important control mechanism in activating ADR1 or NRG1 or both helper NLRs. Future analysis of small molecules and their receptor dynamics can help to develop new strategies for manipulation of plant immunity.

The TIR-catalyzed ADPR-transfer reactions are reminiscent of ADPR chain extensions catalyzed by poly-ADPR polymerases (PARPs) (*36*), which employ a strictly conserved glutamate residue for the catalysis (*36*). The conserved glutamate residue of TIRs may have a similar role in catalyzing ADP-ribosylation of ATP and ADPR by activating the C-1’’ of NAD^+^ in the active site (fig. S9). C-1’’ of ADPr-ATP in the cryo-EM structure retains the same configuration as that of NAD^+^, suggesting that TIR-catalyzed ADP-ribosylation might involve a double-displacement catalytic mechanism as shown for hSARM1 (*34*). However, it is possible that TIR-catalyzed ADPr-ATP or di-ADPR with an inverted configuration at C-1’’ cannot be captured by EDS1-SAG101 due to their deficient or weak EDS1-SAG101 binding activity. Thus, the possibility remains that TIR-catalyzed ADP-ribosylation of ATP and ADPR also involves inversion of the configuration of C-1’’ at this position like PARP-catalyzed ADP-ribosylation of proteins. Regardless of the catalytic mechanism, ADP-ribosylation of ADPR and ATP involves both NADase and PARP-like activities. In addition to ATP and ADPR, ADP can also act as a substrate for a TIR-catalyzed ADP-ribosylation reaction albeit with much less efficiency (Figs. 4A and 5C). This raises the question whether TIRs catalyze ADP-ribosylation of other metabolites. Given the high concentrations of ATP in plant cells, ADPr-ATP is likely the major TIR product for activation of EDS1-SAG101 in the immune response. Biochemical data showed that TIR-mediated hydrolysis of ADPr-ATP and di-ADPR yields pRib-AMP/ADP (Fig. 5C), but it remains possible that TIRs also employ other mechanisms for catalytic production of the two compounds *in vivo*.

## Acknowledgments

We thank the National Protein Science Facility at Tsinghua University for technical assistance, J. Lei, X. Li, X. Fan, and N. Liu at Tsinghua University for data collection, the Tsinghua University Branch of the China National Center for Protein Sciences (Beijing) for providing the cryo-EM facility support and the computational facility support on the cluster of Bio-Computing Platform, Oliver Johanndrees at MPIPZ for providing the EDS1, PAD4 and SAG ortholog sequences. We are grateful to Gordon A Leonard at the ESRF for providing assistance in using beamline ID23-2. A patent application (CN 202210087257.X) has been filed relating to this work.

## Supplementary Materials

### Materials and methods

#### Protein expression and purification in insect cells

For purification of apo-EDS1-SAG101, constructs of full-length *Arabidopsis* EDS1 (residues 1-623, cloned in a pFastbac 1 vector) and full length SAG101 (residues 1-537, cloned in a modified pFastbac 1 vector with a C-terminal twin-StrepII tag) were co-expressed in Sf21 cells at 28 °C for 48 h, using the Bac-to-Bac baculovirus expression system (Invitrogen) protocols. After 48 h of recombinant baculovirus infection, cells were harvested and re-suspended in buffer containing 25 mM Tris (pH 8.0), 150 mM NaCl and 1mM PMSF. The harvested cells were lysed by sonication, the lysate was centrifuged and the supernatant collected. The apo-EDS1-SAG101 complex protein was purified from the supernatant using Streptactin Beads (Novagen). Bound proteins were eluted in a buffer containing 25 mM Tris (pH8.0), 150 mM NaCl, 2.5 mM desthiobiotin and further purified by size exclusion chromatography (Superdex 200 increase 10/300, GE Healthcare) in a buffer containing 25 mM Tris (pH8.0), 150 mM NaCl.

To purify EDS1-SAG101 bound by TIR-generated small molecules, constructs of EDS1 and SAG101 described above were co-expressed with RPP1_WsB (residues 61-1221, cloned in a modified pFastbac 1 vector with a C-terminal twin-StrepII tag) and ATR1_Emoy2 (residues 52-311, cloned in a modified pFastbac 1 vector with a C-terminal 10x His tag) in Sf21 at 28 °C for 48h. Procedures described above were used to purify the EDS1-SAG101 complex protein.

#### Reconstitution of EDS1-SAG101-NRG1A interaction in insect cells

To reconstitute TIR-activated NRG1, N-terminally GST-tagged NRG1A was co-expressed with non-tagged EDS1, C-terminally StrepII-tagged SAG101, RPP1 and ATR1 in Sf21 at 28 °C using Bac-to-Bac baculovirus expression system (Invitrogen) protocols. Insect cells were harvested, lysed and centrifuged. The GST-tagged NRG1A protein was purified using Glutathione Sepharose 4B beads (∼200 µl, Novagen). The resin was washed with 1.0 ml buffer containing 25 mM Tris (pH8.0), 150 mM NaCl for 3 times. Proteins bound to the resin were eluted in buffer containing 25 mM Tris (pH8.0), 150 mM NaCl and 15 mM GSH. The eluted proteins were separated by SDS-PAGE and detected by Coomassie Brilliant Blue staining. Similar protocols were used to reconstitute the EDS1-SAG101-NRG1A interaction with non-tagged RPS4 TIR domain (residues 1-236).

#### Pull-down assays

EDS1-SAG101 protein bound by RPP1 resistosome-catalyzed small molecules was purified and concentrated to 10.0 mg/ml. The concentrated protein (400 µl) in a 1.5-ml centrifuge tube was denatured by heating (92 °C for 7 min). The sample was centrifuged to remove precipitate and the supernatant was collected to repeat the heating-centrifugation steps two more times. The final 400 µl supernatant was divided equally in two 1.5-ml centrifuge tubes. N-terminally GST-tagged ADR1-L1/NRG1 was expressed in Sf21 insect cells and purified using GS4B beads. GS4B beads bound by GST-tagged ADR1-L1/NRG1 were incubated with excess apo-EDS1-PAD4/ apo-EDS1-SAG101 and 200 µl supernatant obtained above on ice for 30 min. The GS4B beads were then washed three times with 1.0 ml buffer containing 25 mM Tris (pH 8.0), 150 mM NaCl. Proteins bound to the resins were eluted with buffer containing 25 mM Tris (pH 8.0), 150 mM NaCl and 15 mM GSH, analyzed by SDS-PAGE and detected by Coomassie Brilliant Blue staining.

#### Chemical synthesis of di-ADPR

Compound **5** was synthesized from compound **1** in four steps *via* protection of the 2*^’^*-and 3*^’^*-OH group and subsequent phosphoramidation of the 5*^’^*-OH group (Data S1, Scheme 1). Similarly, starting from copound **6** and following a sequence of protection, lactone reduction, protection, deprotection, and final phosphoramidite installation furnished compound **12** (Data S1, Scheme 2). The absolute configuration of compound **12** is confirmed by NOE correlations.

The synthesis of the dimeric ADP-Ribose **32** commenced with *N*-benzoyladenosine **13** (Data S1, Scheme 3). Selective silylation of nucleoside **13** followed by condensation with beta-D-ribofuranose 1-acetate 2,3,5-tribenzoate gave intermediate **15**. A series of protecting group manipulation resulted in the formation of compound **19**, which was phosphorylated to provide **20**. Following similar sequence of transformations, the second phosphate group was installed to provide **25**. Selective deprotection of **25** produced compound **26** which reacted with compound **5** to give compound **27**. Reveal of the phosphoric acid functionality **29** followed by reaction with **12** generated compound **30**. Finally, global deprotection provided the final product **32**.

#### LC-HRMS assay

Purified apo-EDS1-SAG101 proteins (0.3 mg) were dissolved in 20 µl of 0.1M NH_4_HCO_3_ buffer (PH 7.6). Four volumes of methanol were added to precipitate proteins and extract small molecules. The mixed solution was centrifuged at 14,000 g to remove the precipitate after stored in a −80°C fridge for 30 min. The supernatant was transferred to a new 1.5-ml centrifuge tube prior to mass spectrometry analysis. An orbitrap mass spectrometer (Q-Exactive HFX, Thermo Fisher, CA) coupled with a Vanquish UHPLC system (Thermo Fisher, CA) was applied to analyze small molecules. A BEH Amide column (100 × 2.1 mm, Waters, USA) was used for LC separation with injection volume of 5 µl. Mobile phase A and B contained 95% acetonitrile and aqueous with 5 mM ammonium acetate respectively. The gradient of 10% to 50% mobile phase B within 5 min was applied. Data dependent MS/MS acquisition (DDA) with full scan of m/z 200-1500 in negative ion mode was performed. Mass resolutions of MS and MS/MS scans were set as 60,000 and 15,000, respectively. The source parameters were set as follows: spray voltage: 3 kv; capillary temperature: 320 °C; heater temperature: 300 °C; sheath gas flow rate: 35 arb; auxiliary gas flow rate: 10 arb. Xcalibur 4.4 (Thermo Fisher, CA) was utilized for data analysis.

### TIR domain *in vitro* enzyme activity assay

The RPS4 TIR domain (residues 1-236) was cloned in a pET30a vector and expressed in *Escherichia coli* BL21(DE3) stains at 16°C with 0.6 mM IPTG for 16 h. The cells were harvested, re-suspended and lysed in buffer containing 25 mM Tris (pH 8.0) and 150 mM NaCl. After sonication and centrifugation, RPS4 TIR domain protein was purified using Ni-NTA (Novagen). After washing Ni-NTA beads (∼2 ml) three times with 7.0 ml buffer containing 25 mM Tris (pH 8.0), 150 mM NaCl and 15 mM imidazole for 3 times, 1 ml ddH_2_O containing 1 mM NAD^+^, 1 mM NAD^+^ + 1 mM ADP, 1 mM NAD^+^ + 1 mM ATP was added to the beads. The mixtures were left at room temperature for 16 h. 200 µl from each reaction was used to test EDS1-SAG101 interaction with NRG1A as described in ‘Pull-down assay’.

LC-HRMS was used to further identify produced small molecules. ∼0.5 mg apo-ESD1-PAD4 or apo-EDS1-SAG101 protein were incubated with above 1 ml ddH_2_O solution after reaction on ice for 2 h. Size exclusion chromatography (Superdex 200 increase 10/300, GE Healthcare) was used to replace ddH_2_O solution buffer to 0.1M NH_4_HCO_3_ buffer (pH 7.6). The protein was concentrated to 20 µl, and 80 µl methanol added to denature the protein. The mixture was stored at −80°C for 30 min and then centrifuged to remove precipitate. The supernatant was used for LC-HRMS.

### Cryo-EM sample preparation and data acquisition

EDS1-SAG101 purified proteins were concentrated to ∼1mg/ml. Aliquots of prepared proteins were applied to holey carbon grids (Quantifoil Au R1.2/1.3 300 mesh) glow-discharged for 30 s at medium level in Harrick Plasma after 2 min evacuation. Then, the grids were blotted for 3 s at 100% humidity at 8°C, frozen and plunged into liquid ethane cooled with liquid nitrogen with Vitrobot Mark IV (Thermo Fisher Scientific).

Data were collected in counted super-resolution mode on a Titan Krios3 (FEI) operating at 300 kV with a BioQuantum imaging filter (Gatan) and K3 direct detection camera (Gatan) at ×81,000 magnification, with a physical pixel size of 0.54125 Å using AutoEMation (*38*). Each stack was exposed for 2.56 s with an exposure time of 0.08 s per frame and recorded as a movie of 32 frames, resulting in the total dose rate of approximately 50 electrons/Å^2^ for each movie stack. Details could be found in Table S1.

### Image processing and 3D reconstruction

The stacks were motion-corrected with MotionCor2 and then 2 × 2 binned (*39*), resulting in a pixel size of 1.0825 Å per pixel (*39*). The average of each movie stack was calculated and summed, then the contrast transfer function (CTF) parameters were estimated by CTFFIND4 program and dose-weighted average for reconstruction with RELION-3.1 (*40–43*).

The cryo-EM processing workflow for EDS1-SAG101 was outlined in fig. S2. Total of 4,860 micrographs were collected and 6,993,337 particles were autopicked in RELION-3.1 using Laplacian-of-Gaussian. Then two rounds of reference-free 2D classification (150 classes and 30 iterations each time) were performed to remove contaminants and noise in the raw automatically picked particles. After 2D classification, the selected particles were subjected to ab-initio reconstruction requesting three classes in RELION-3.1. The best 3D map was subjected to initial 3D classification. 785,900 particles were selected for the second round of 3D classification. Last, a total of 652,337 particles were selected and subjected to a 3D auto-refinement with an overall mask. After CTF refinement and Postprocessing, yielding a density map at a resolution of 2.71 Å, based on the gold standard FSC 0.143 criteria (*44*). Local resolution distribution was evaluated using RELION (*45*).

### Cryo-EM model building and refinement

The EM density reconstructed from EDS1-SAG101 was used for initial model building. The crystal structure of apo-EDS1-SAG101 (PDB 4NFU) was docked into the EM density in Chimera to build a model of EDS1 (*15, 46*). Manual atomic model building was carried out in the software Coot using sequences of SAG101 (*47*). An ADPr-ATP molecule was readily discernible in the EM density and was included in the model between EDS1 and SAG101. Phenix.real_space.refine was used to refine the model of EDS1-SAG101 with the final EM map. The refined models were validated using phenix.validation_cryoem (*48*). Model validations were performed using MorProbity and EMRinger included in the PHENIX package (*48*). Model statistics can be found in Table S1.

### Purification, crystallization of RPP1^TIR^ and data collection

RPP1^TIR^ (residue 83-254) was cloned into pFastBac1 (Invitrogen) with an N-terminal GST tag. The construct was used for generating recombinant baculovirus in sf21 insect cells (Invitrogen). RPP1^TIR^ was expressed in sf21 insect cells with recombinant baculovirus infection at 28 °C for 60h. The infected cells were harvested and lysed by sonification in buffer containing 25 mM Tris-HCl pH 8.0, 150 mM NaCl. The cell lysates were centrifuged at 30,000 *g* for 1.5 h. The supernatant containing soluble proteins were collected and allowed to flow through GST4B resin (GE Healthcare). After washing with three column volumes of sonification buffer, the fusion proteins were incubated with PreScission protease at 4°C overnight to remove the N-terminal GST tag. The digested RPP1^TIR^ proteins flowed through the columns in the buffer containing 25 mM Tris-HCl pH 8.0, and 150 mM NaCl.

Purified RPP1^TIR^ proteins were concentrated to 8 mg/ml after size exclusion chromatography (SEC). RPP1^TIR^ (residue 83-254) was incubated with 4 mM NAD^+^. After 2 h, the RPP1^TIR^-NAD^+^ mixture was centrifuged at 12,000 *g* for 10 min and immediately applied for crystal screening. Protein crystals were generated by mixing the protein with an equal amount of well solution (1.0 μl) by the hanging-drop vapor-diffusion method. After several rounds of crystallization condition optimizations, the best RPP1^TIR^ (residue 83-254) crystals were obtained under the condition of 20 % w/v polyethylene glycol 3000 (PEG3000), 100 mM HEPES pH 7.5, 200 mM sodium acetate. The crystals were flash frozen in liquid nitrogen using 12.5 % w/v glycerol as the cryoprotecting buffer and sent for synchrotron in Grenoble, France.

### Crystal structure determination and refinement of RPP1^TIR^

RPP1^TIR^ diffraction data was collected at the ID23-2 at the European Synchrotron Radiation Facility (ESRF), Grenoble, France. The data was processed using XDS and Pointless (1+2) (*49,50*). The crystal structure of RPP1^TIR^ was determined by molecular replacement (MR) with PHASER (51) using structure of RPP1^TIR^ from the RPP1 resistosome (PDB code: 7CRC) as the initial searching model. The model from MR was built with the COOT (*47*) and subsequently subjected to refinement by Phenix_real_space_refine (*48*). Statistics of diffraction data and refinement of the EDS1-PAD4 model are summarized in Table S1. Structural figures were prepared using the program PyMOL (http://www.pymol.org/).

### Semi-*in vitro* ADPr-ATP induced EDS1-SAG101-NRG1 association analysis

EDS1-FLAG, SAG101-FLAG and NRG1A^E14A/E27A^-SH were co-expressed in *Nb-epss*. Nine 1 cm leaf discs were collected at 48 hpi. Total protein from the plant material ground to fine powder was extracted with 2 ml extraction buffer (100 mM Tris-HCl (pH 8.0), 150 mM NaCl, 1 mM EDTA, 0.5% NP40 (v/v), 10 mM DTT, 1x Plant protease cocktail (11873580001, MilliporeSigma)). Tubes were vortexed, then samples left on ice for 30 min and mixed several times. Samples were then centrifuged for 10 min at 16500 × *g*. The supernatant was transferred to 2 ml Eppendorf tubes and 50 µl aliquots were taken as input samples. 1.8 ml supernatant was divided into two 1.5 ml protein low-binding Eppendorf tubes. 80 µl 50% ethanol was added to 80 µl enriched EDS1-SAG101 protein bound by RPP1 resistosome-catalyzed ADPr-ATP to denature proteins. The mixture was stored at −20 degree for 20 min and centrifuged for 15 min at 16,500 × g. 7.5 µl supernatant including ADPr-ATP and 25% ethanol was added separately into two samples and tubesrotated for 1 h at 4 °C. 1.5 µl ADPr-ATP and 25% ethanol was added to samples after 1 h incubation, and co-IPs were conducted for 2 h with 15 μl α-HA affinity matrix (11815016001, MilliporeSigma) under constant rotation. Beads were collected by centrifugation at 1,200 × g for 2 min and washed three times in extraction buffer (without DTT). All co-IP steps were conducted at 4 °C. Beads and input samples were boiled at 95 °C in 60 μl 2 × Laemmli buffer and proteins separated by SDS-PAGE followed by immunoblotting. Antibodies used for immunoblotting were α-HA (1:5000; MMS-101P, BioLegend), α-FLAG (1:5000; f7425, MilliporeSigma), Goat anti-Rb IgG secondary HRP conjugated antibody (1:5000; 31460, invitrogen), Goat Anti-Rat IgG secondary HRP conjugated antibody (1:5000; AP136P, Sigma-Aldrich).

---

**Fig. S1.**
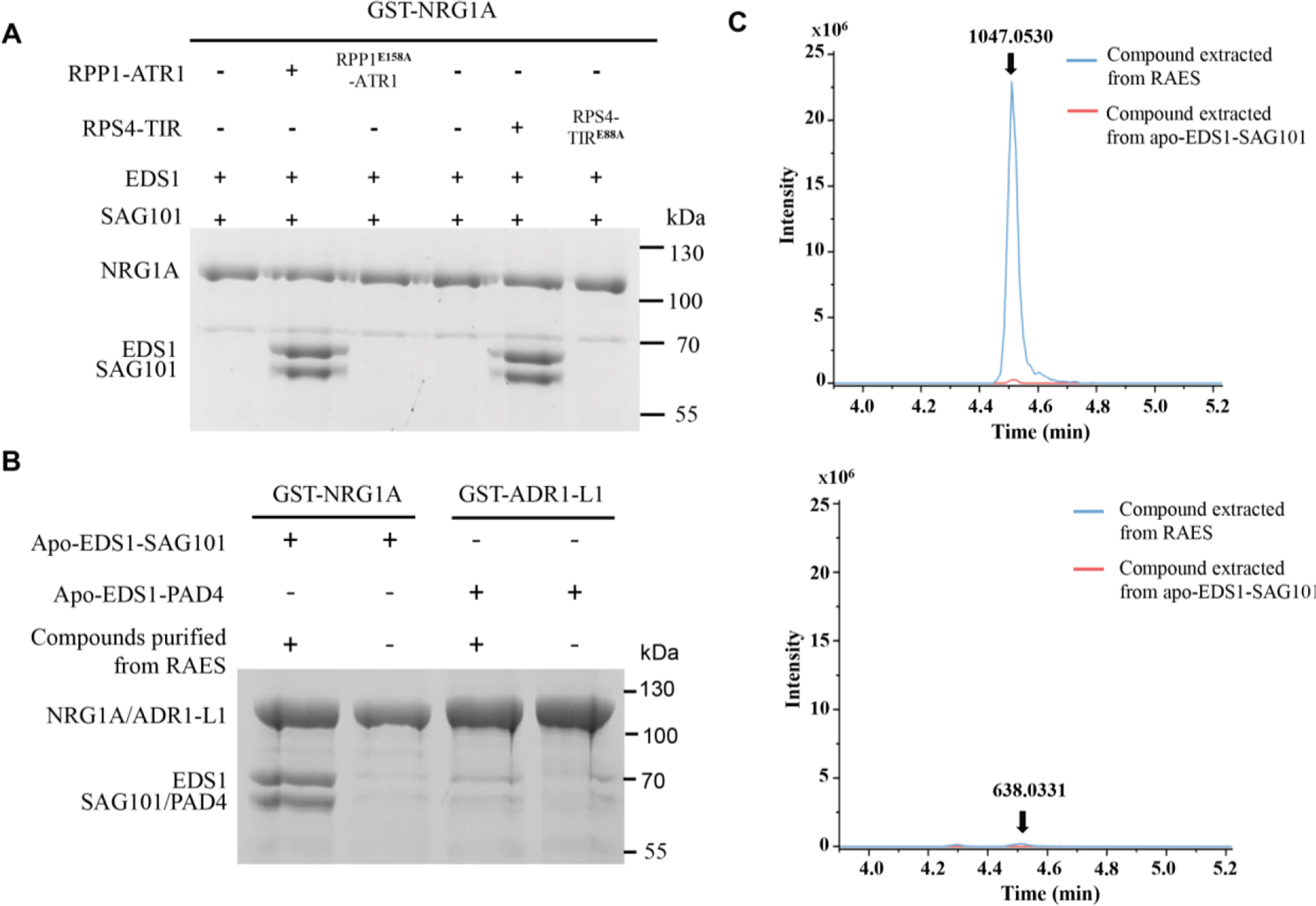
TIR-catalyzed products induce EDS1-SAG101 interaction with NRG1A in insect cells. (A) RPP1-ATR1 induces EDS1-SAG101 interaction with NRG1A in insect cells. N-terminally GST-tagged NRG1A was co-expressed with non-tagged EDS1, C-terminally strep-tagged SAG101, RPP1 (wild type or catalytic mutant E158A) and 10xHis-tagged ATR1 or RPS4-TIR (residues 1-236, wild type or catalytic mutant E88A) in insect cells. GST-NRG1A was purified using Glutathione Sepharose 4B (GS4B) beads. Bound proteins were eluted, separated by SDS-PAGE and detected by Coomassie Brilliant Blue staining. (**B**) EDS1-SAG101 was expressed as described in (A) and purified with Strep-Tactin resin. The purified protein was concentrated and denatured by heating. After centrifugation, the supernatant was collected and incubated with purified N-terminally GST-tagged NRG1A/ADR1-L1 and apo-EDS1-SAG101/apo-EDS1-PAD4. The mixture was flowed through GS4B beads and bound proteins were eluted, analyzed by SDS-PAGE and detected by Coomassie Brilliant Blue staining. (**C**) Chromatograms of supernatant extracts from the denatured apo-EDS1-SAG101 (red) and EDS1-SAG101 co-expressed with RPP1-ATR1 (blue). Purified apo-EDS1-SAG101 and EDS1-SAG101 protein denatured with 80% methanol (v/v) and subjected to LC-HRMS analyses. The peak of the top LC was corresponding to the ion mass of 1047.0530 (-). The peak of the bottom LC was corresponding to the ion mass of 638.0331 (-).

**Fig. S2.**
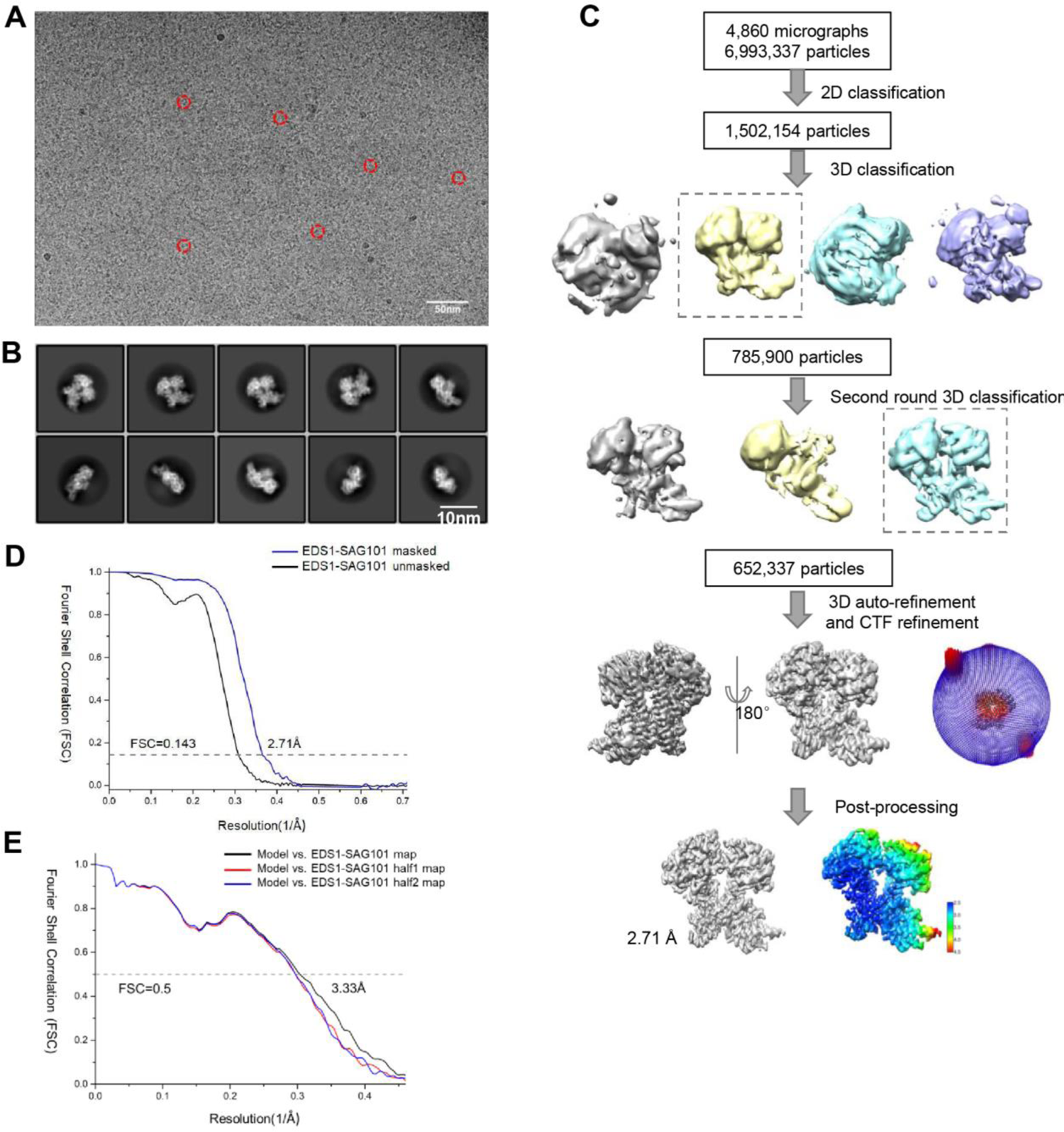
Cryo-EM reconstruction of the EDS1-SAG101 complex. (**A**) Representative cryo-EM micrograph of the EDS1-SAG101 complex. (**B**) Representative views of 2D class averages of the EDS1-SAG101 complex. (**C**) The cryo-EM image processing workflow. (**D**) FSC curves at 0.143 of the final reconstruction of the EDS1-SAG101 complex unmasked (black) or masked (blue). (**E**) FSC curves at 0.5 for model refined against the final map (black), the first half map (red) and the second half map (blue).

**Fig. S3.**
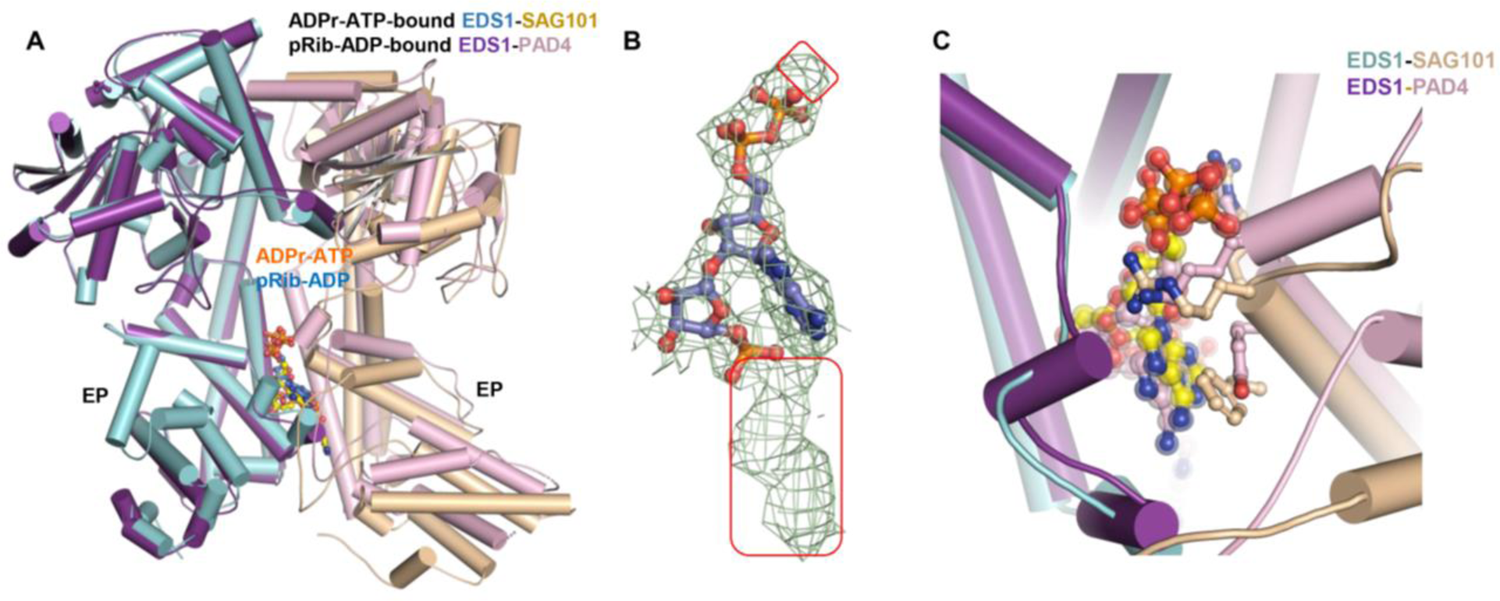
ADPr-ATP and pRib-ADP binding a conserved site. (**A**) Structural alignment between ADPr-ATP-bound EDS1-SAG101 and pRib-ADP-bound EDS1-PAD4. (**B**) Modeling of pRib-ADP into the cryo-EM density between EDS1 and SAG101 EP domains that is not occupied by EDS1-SAG101. Cryo-EM densities not occupied by the modeled pRib-ADP are highlighted with open frames. (**C**) Structural alignment between ADPr-ATP-bound EDS1-SAG101 and pRib-ADP-bound EDS1-PAD4 around the ligand-binding site. Residues from SAG101 and PAD4 interacting with ADPr-ATP and pRib-ADP, respectively, are shown in stick.

**Fig. S4.**
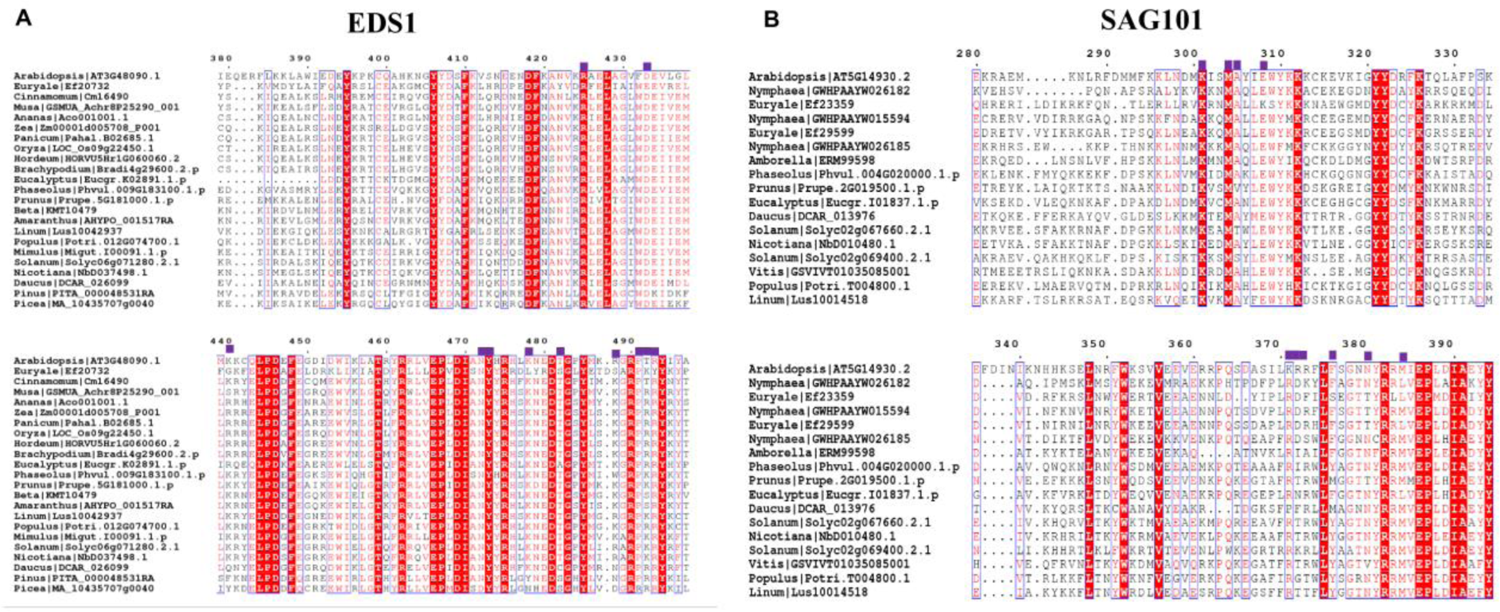
Sequence alignment of EDS1 and SAG101 from different plant species. **(A**) Sequence alignment of EDS1 proteins from different plant species. Amino acids involved in interaction with ADPr-ATP are highlighted with purple squares on the top. (**B**) Sequence alignment of SAG101 proteins from different plant species. Amino acids involved in interaction with ADPr-ATP are highlighted with purple squares on the top. EDS1 and SAG101 sequences were predicted using the EP-domain HMM (PFAM: PF18117) and classified using a ML phylogenetic tree (*52*).

**Fig. S5.**
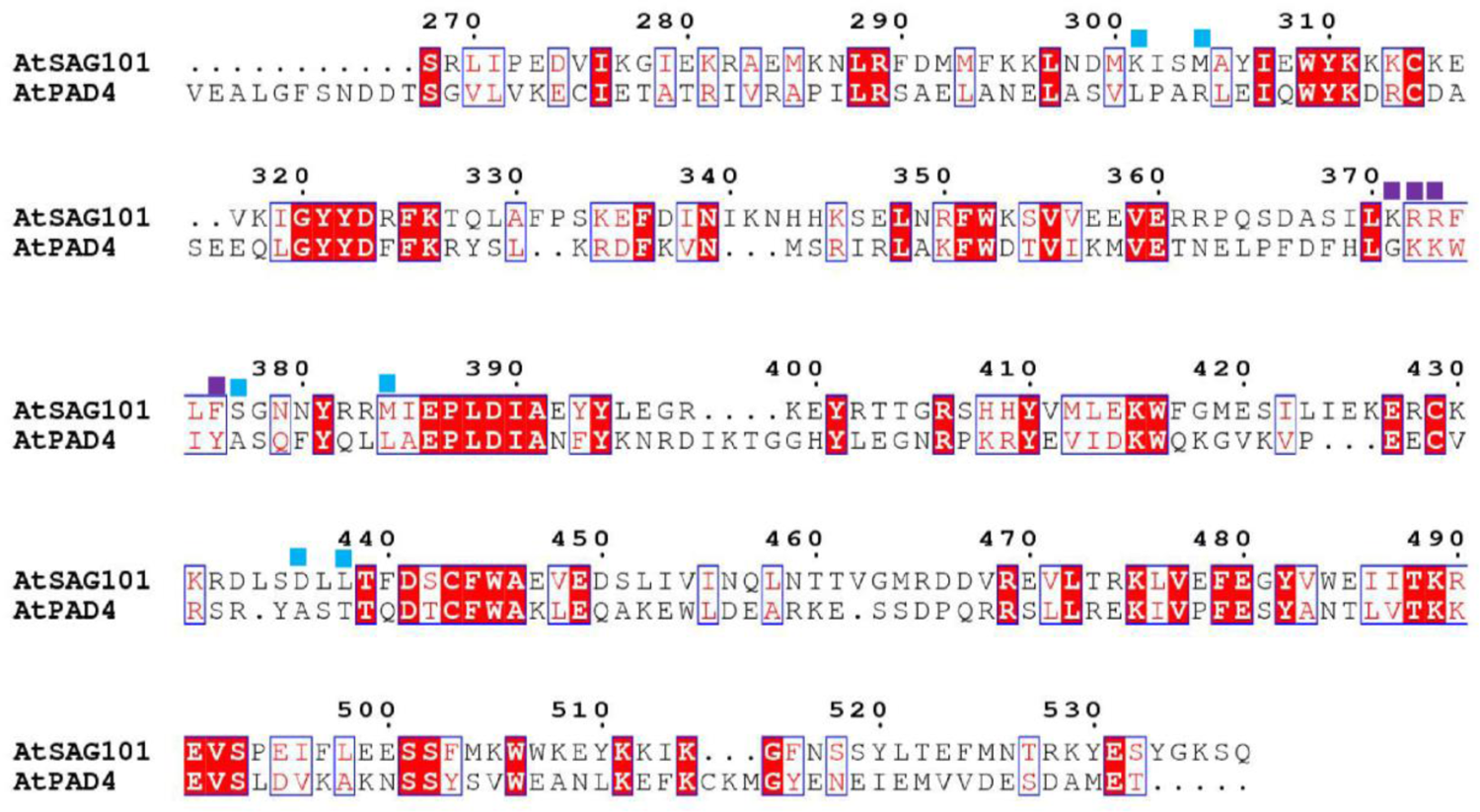
Sequence alignment of SAG101 and PAD4 in *Arabidopsis* Amino acids of SAG101 involved in interaction with the pRib-ADP and ADPR moieties of ADPr-ATP are highlighted with purple and cyan squares on the top, respectively.

**Fig. S6.**
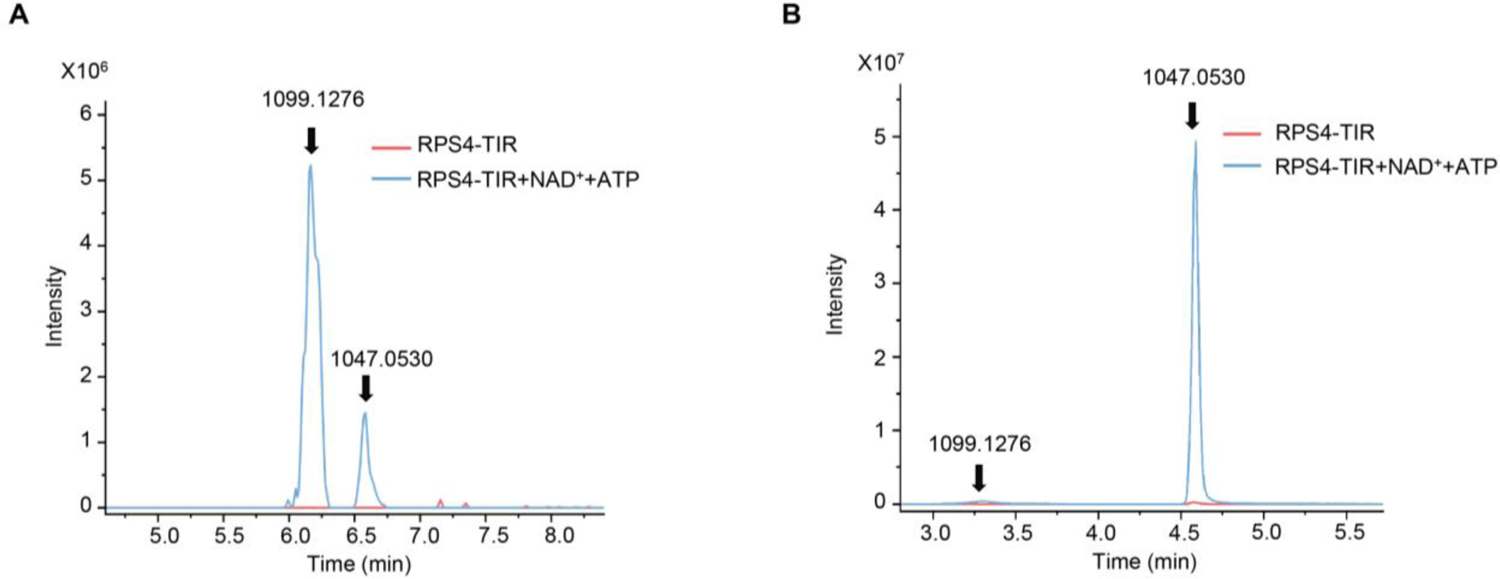
TIR-catalyzed production of di-ADPR and ADPr-ATP. (**A**) LC-HRMS chromatograms of small molecules from RPS4-TIR catalyzed products with NAD^+^ + ATP as substrates. (B) LC-HRMS chromatograms of EDS1-SAG101-captured small molecules from RPS4-TIR catalyzed products with NAD^+^ + ATP as substrates. The assays were performed as described in Figure 4B.

**Fig. S7.**
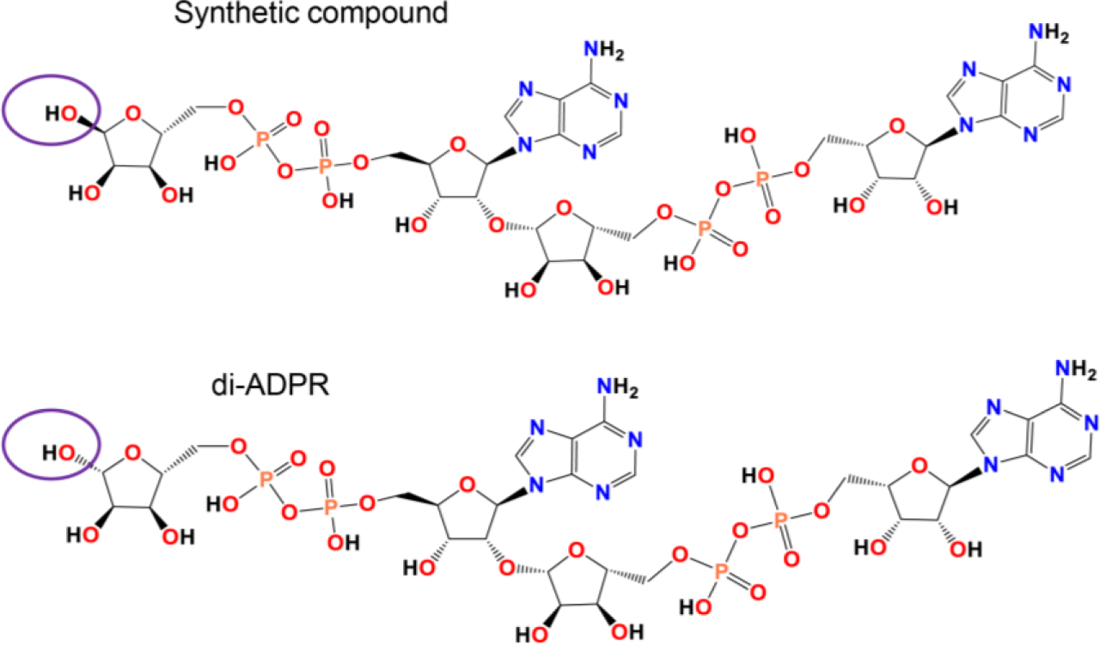
Structural comparison of the synthetic di-ADPR enanthiomer and di-ADPR. Molecular structures of the synthetic di-ADPR enanthiomer (top) and di-ADPR (bottom). The difference between the compound is highlighted within the elliptical frames.

**Fig. S8.**
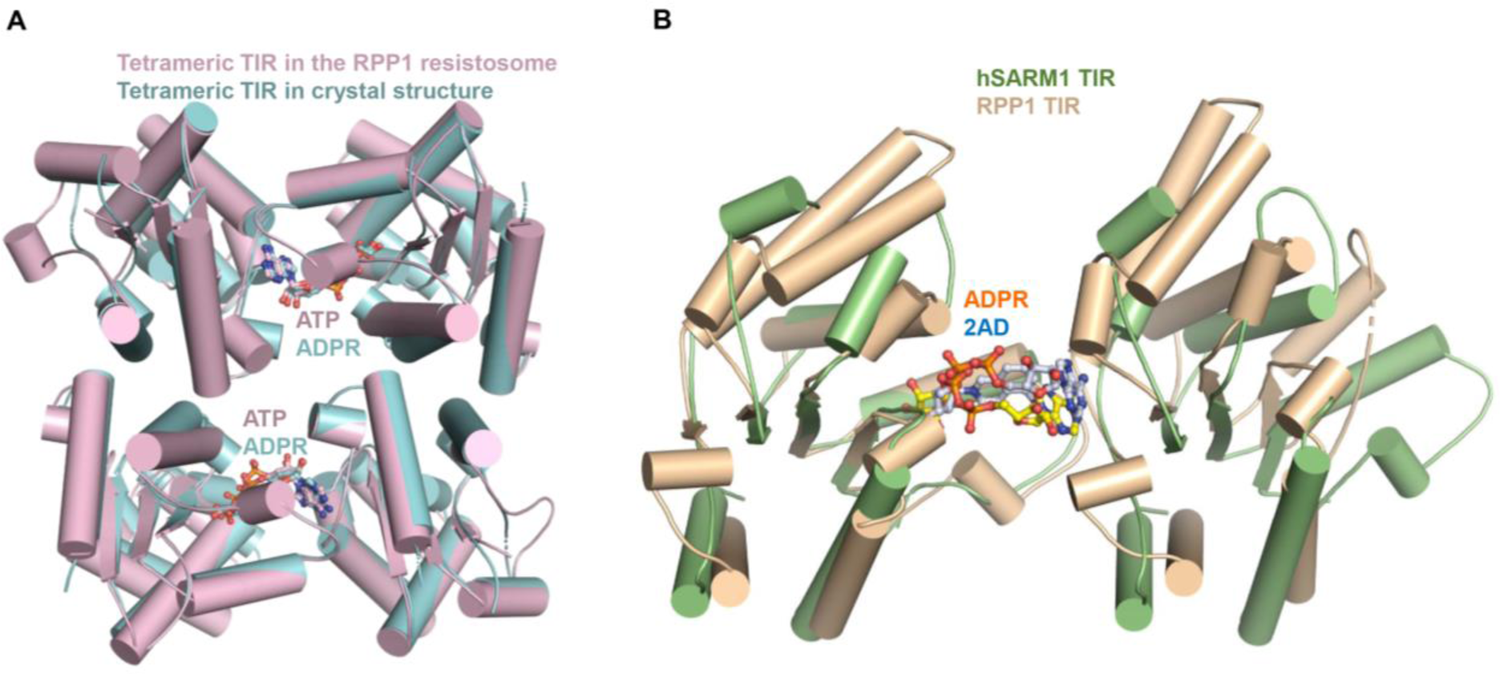
Structural comparison of RPP1 crystal structure with RPP1 TIR from the RPP1 resistosome and hSARM1 TIR. (**A**) Crystal structure of tetrameric RPP1 TIR was aligned with the four TIR domains from the cryo-EM structure of the RPP1 resistosome (PDB code: 7CRC). (**B**) Structural superposition of an ADPR-bound asymmetric RPP1 TIR dimer and 2AD (inhibitor of hSARM1)-bound asymmetric hSARM1 TIR dimer (PDB code: 7NAH)

**Fig. S9.**
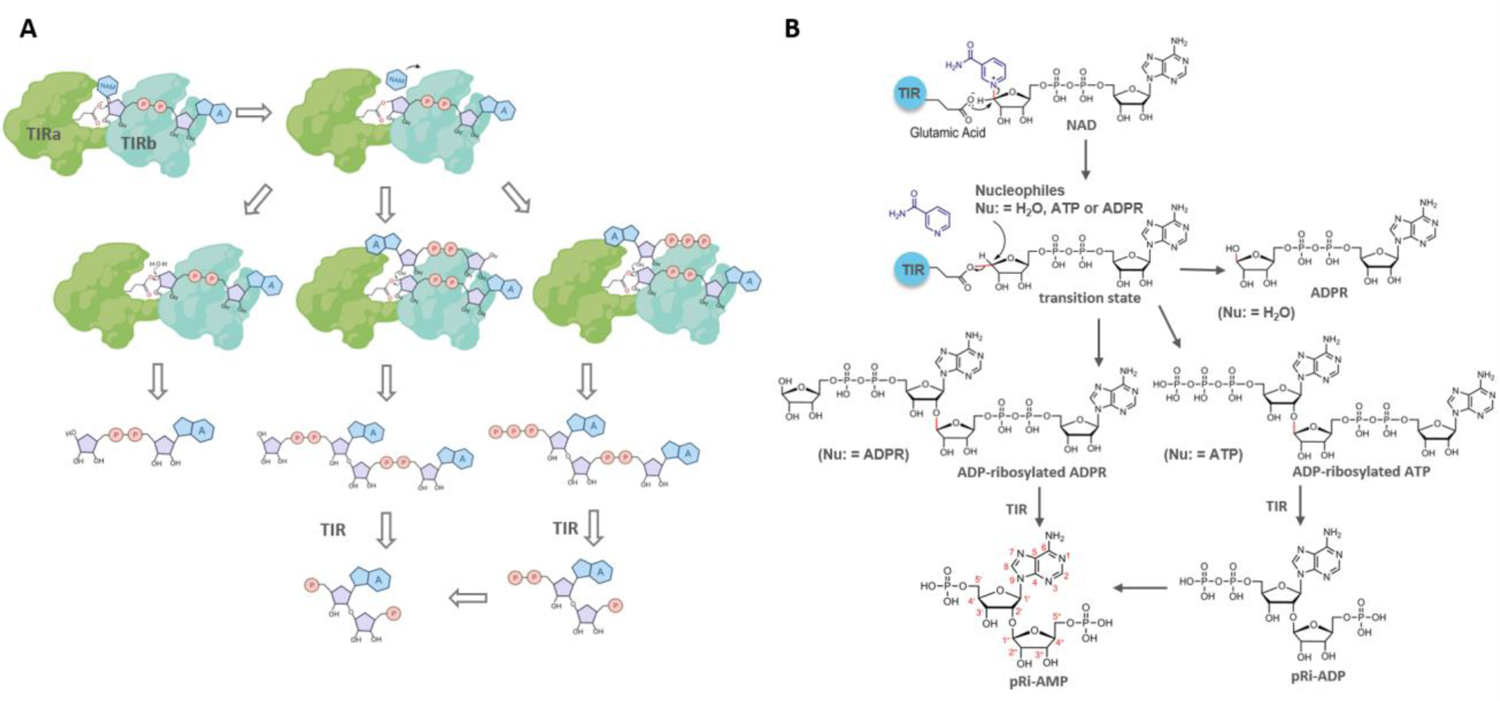
Model on TIR-catalyzed ADP-ribosylation reaction. (**A**) A cartoon illustration of proposed mechanism for TIR-catalyzed production of ADPR, di-ADPR, ADPr-ATP, pRib-AMP and pRib-ADP. Asymmetric homodimer TIR proteins forming a basic catalytic unit are colored in green. At the active site, the catalytic Glutamic acid is shown. Sugar rings are represented by grey pentagons and phosphate groups by circled letter P. NAM and A represent Nicotinamide and Adenine, respectively. (**B**) Catalytic mechanism in (A) presented in molecular structures. NAD^+^ as a substrate binds to the active site formed by an asymmetric TIR dimer. The positively charged NAM group of NAD^+^ as a leaving group is replaced by the catalytically active glutamic acid probably via an S_N_2 substitution reaction, resulting in a transition state that is chemically suceptible to nucleophiles such as H_2_O, ATP and ADPR. The following substitution of glutamic acid by H_2_O, 2′-OH of ADPR and ATP produces ADPR, ADP-ribosylated ADPR (di-ADPR) and ADP-ribosylated ATP (ADPr-ATP), respectively. The latter two products can be further hydrolyzed to pRib-AMP and pRib-ADP by TIR proteins. In addition, pRib-ADP might be converted into pRib-AMP through hydrolysis. The chemical bonds involved in the reactions are indicated by red lines. Atom numbering of chemical nomenclature is shown in red in pRib-AMP.

**Table S1.** Cryo-EM data collection, refinement and validation

|  |  |
| --- | --- |
| PDB and EMD ID | EDS1-SAG101<br>7XJP EMD-33233 |
| **Data collection** |  |
| Cryo electron microscope | FEI Titan Krios |
| Voltage (kV) | 300 |
| Detector | Gatan K3 Summit |
| Energy filter slit width (eV) | 20 |
| Magnification | 81,000 x |
| Pixel size (Å) | 1.0825 |
| Total electron exposure (e<sup>-</sup>/Å<sup>2</sup>) | 50 |
| Number of frames collected | 32 |
| Defocus range (μm) | -1.3 ~ -1.8 |
| Automation software | AutoEMation |
| Micrographs collected | 4,965 |
| Micrographs used | 4,860 |
| **3D reconstruction** |  |
| Software | RELION 3.1 |
| Total extracted particles | 6,993,337 |
| Total number particles for final refinement | 652,337 |
| Symmetry imposed | C1 |
| Resolution range (Å) | 2.53~4.38 |
| Resolution (Å) after refinement (FSC=0.143) | 3.25 |
| Resolution (Å) after post-processing (FSC=0.143) | 2.71 |
| Map sharpening B-factor (Å<sup>2</sup>) | -30 |
| **Refinement and validation** |  |
| Software | Phenix.real_space_refine |
| Model resolution (Å) | 3.1(FSC=0.5) |
| Model composition |  |
| Non-hydrogen atoms | 8930 |
| Protein residues | 1089 |
| Map-model CC (overall/local) | 0.71/0.69 |
| B factors (Å<sup>2</sup>) | 118.6 |
| R.M.S deviations |  |
| Bonds lengths (Å) | 0.003 |
| Bonds angles (°) | 0.690 |
| MolProbity score | 1.73 |
| Clash score | 11.44 |
| Rotamer outliers (%) | 1.02 |
| Cb outliers (%) | 0.00 |
| CaBLAM outliers (%) | 0.84 |
| EMRinger score | 2.56 |
| Ramachandran plot statistics |  |
| Preferred (%) | 97.22 |
| Allowed (%) | 2.78 |
| Outlier (%) | 0.00 |

**Table S2.** Data collection and refinement statistics

| Data collection |  |
| --- | --- |
| Data set | RPP1<sup>TIR</sup> |
| PDB ID | 7XOZ |
| Wavelength (Å) | 1.000 |
| Resolution (Å) | 39.12-2.52 (2.61-2.52) |
| Space group | P 2<sub>1</sub> |
| Cell dimensions | 69.80 Å, 81.13 Å, 70.71 Å, 90.00°, 112.33°, 90.00° |
| Unique reflections | 24611 (2370) |
| Completeness (%) | 98.43 (90.79) |
| R-meas (%) | 29.32 (199.1) |
| R-pim | 13.7 (92.96) |
| Redundancy | 4.3 (4.3) |
| Average I/σ(I) | 3.97 (0.52) |
| Statistics for Refinement |  |
| Rwork (%) | 25.8 (38.8) |
| Rfree (%) | 29.9 (41.3) |
| Reflections used in refinement | 24381 (2239) |
| Number of non-hydrogen atoms | 5288 |
| ligands | 120 |
| solvent | 42 |
| Protein residues | 645 |
| R.m.s.d. |  |
| Bond (degree) | 0.92 |
| Length (Å) | 0.009 |
| Ramachandran plot |  |
| Favored region (%) | 96.84 |
| Allowed region (%) | 2.69 |
| Outliers (%) | 0.47 |
| Average B-factor | 61.14 |
| ligands | 54.64 |
| solvent | 54.82 |

**Table S3.**
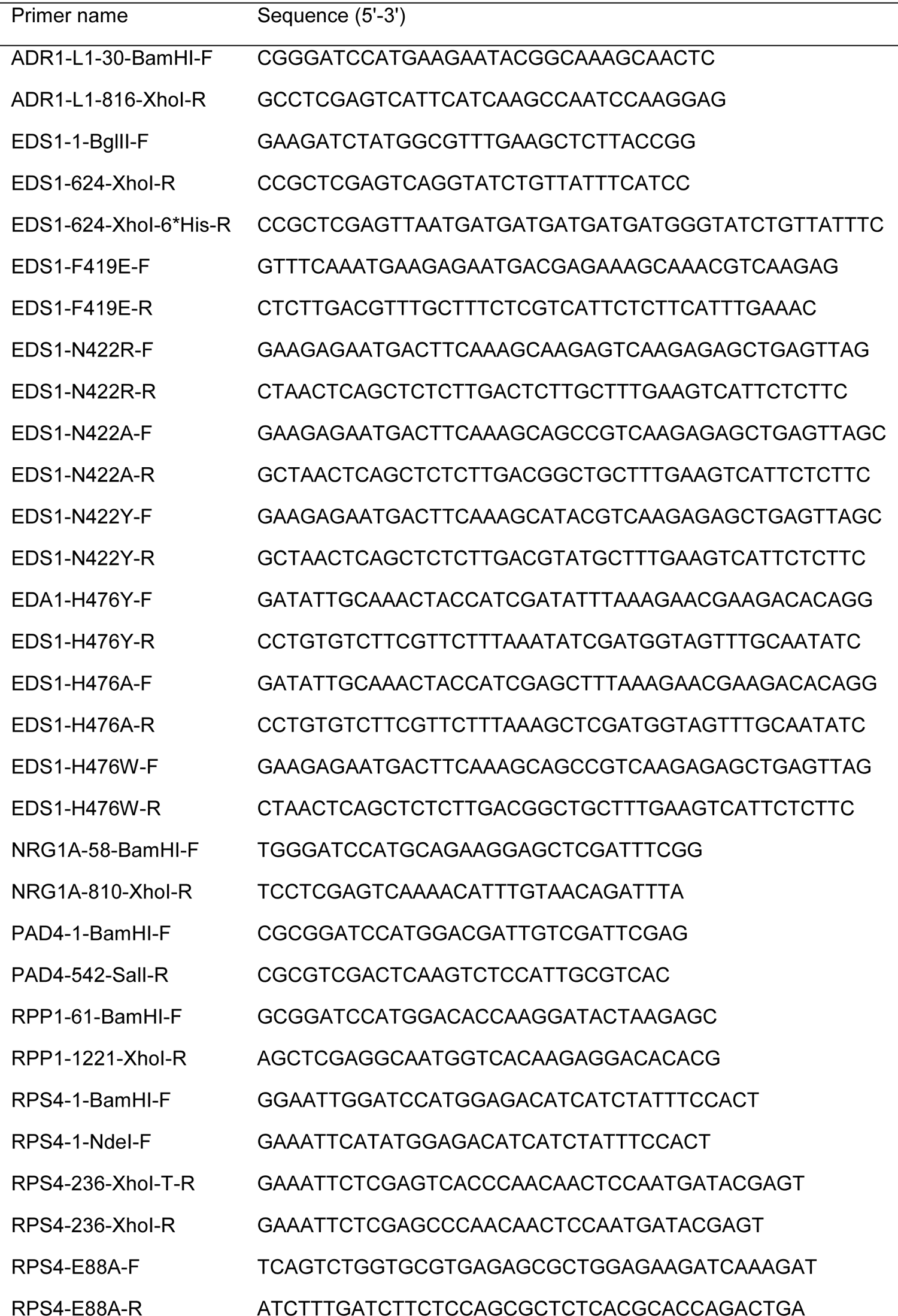
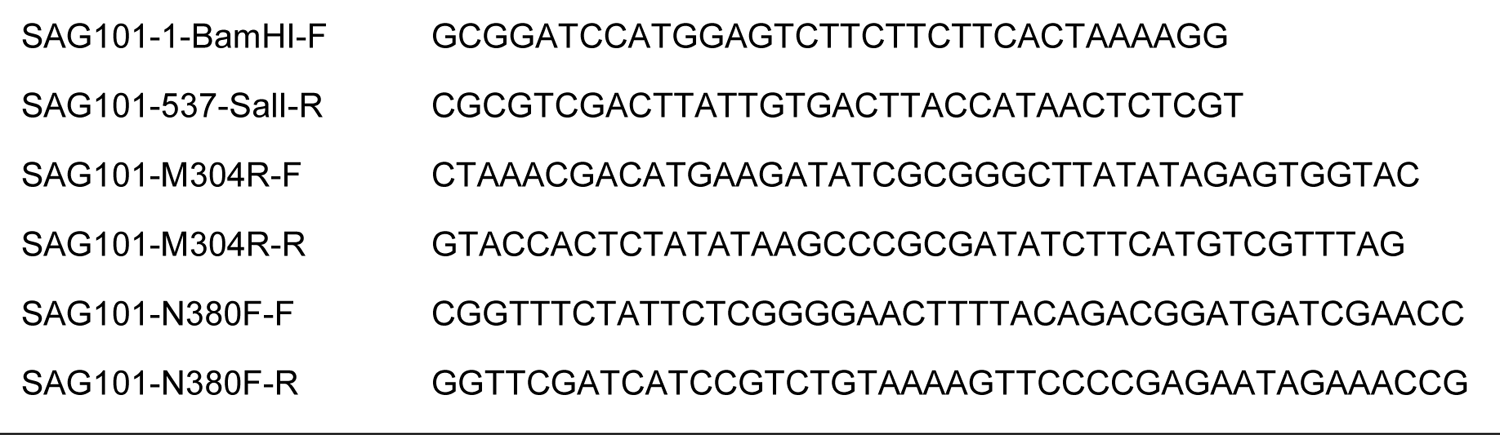
Oligonucleotides used in this study

